# An algorithm for automated, noninvasive detection of cortical spreading depolarizations based on EEG simulations

**DOI:** 10.1101/393058

**Authors:** Alireza Chamanzar, Shilpa George, Praveen Venkatesh, Maysamreza Chamanzar, Lori Shutter, Jonathan Elmer, Pulkit Grover

## Abstract

We present a novel signal processing algorithm for automated, noninvasive detection of Cortical Spreading Depolarizations (CSDs) using electroencephalography (EEG) signals and validate the algorithm on simulated EEG signals. CSDs are waves of neurochemical changes that suppress neuronal activity as they propagate across the brain’s cortical surface. CSDs are believed to mediate secondary brain damage after brain trauma and cerebrovascular diseases like stroke. We address key challenges in detecting CSDs from EEG signals: (i) decay of high spatial frequencies as they travel from the cortical surface to the scalp surface; and (ii) presence of sulci and gyri, which makes it difficult to track the CSD waves as they travel across the cortex. Our algorithm detects and tracks “wavefronts” of the CSD wave, and stitches together data across space and time to decide on the presence of a CSD wave. To test our algorithm, we provide different models and complex patterns of CSD waves, including different widths of CSD suppressions, and use these models to simulate scalp EEG signals using head models of 4 subjects from the OASIS dataset. Our results suggest that the average width of suppression that a low-density EEG grid of 40 electrodes can detect is 1.1 cm, which includes a vast majority of CSD suppressions, but not all. A higher density EEG grid having 340 electrodes can detect complex CSD patterns as thin as 0.43 cm (less than minimum widths reported in prior works), among which single-gyrus propagation is the hardest to detect because of its small suppression area.

## I. INTRODUCTION

IN this work, we provide an algorithm for noninvasive and automated detection of Cortical Spreading Depolarizations (CSDs) which is tested on simulated electroencephalography (EEG) signals. CSDs are waves of neurochemical changes that propagate slowly (1 to 8 mm/min) across the cortical surface and result in a suppression of normal neuronal electrical activity [2]–[4]. CSDs occur because of loss of electrochemical gradient across neuronal membranes [5]. Increasing evidence shows that CSDs may contribute to secondary brain injury after trauma, stroke, and hemorrhage by causing microvascular constriction and brain tissue hypoperfusion [5]–[7]. Approximately 2.5 million TBIs occur in the United States per year^1^, and among all of the deaths due to injuries, 30% are due to TBIs^2^ [8]. Higher frequency of occurrence of CSDs is correlated with higher tissue damage [6], and surgical measures that reduce CSD incident lead to improved clinical outcomes [9]. Therefore, there is a need for early detection and inhibition of CSDs. At present, aside from visual examination of electrophysiological data [10], there is no established technique for automated noninvasive detection of CSDs. We view our work as the first step in that direction.

**Fig. 1:**
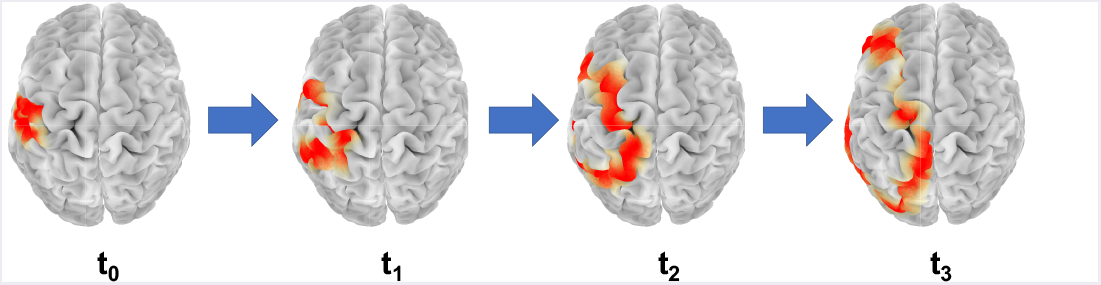
A simulation of propagation of a CSD wave on the cerebral cortex across the left hemisphere over time *t*_0_ < *t*_1_ < *t*_2_ < *t*_3_. The red region in the figure is the region of suppression of normal brain activity.

Two main challenges in detecting CSD from noninvasive EEG recording are: (i) transient disappearance of parts of a CSD wave from EEG recordings when it enters sulci; and (ii) spatially blurred observation of the underlying cortical activity recorded by scalp EEGs. This spatial blurring is due to the decay of high resolution information passing through bone and soft tissue, which complicates detection of narrow CSDs. In idealized spherical models, a CSD wave forms an annular ring of depressed brain activity that propagates across the cortical surface. However, due to the presence of folds (gyri and sulci) in the human cortex, a CSD wave would appear in EEG recordings as broken parts that we call “wavefronts,” which can evolve to break down or combine with author-notes wavefronts as the wave propagates. To address the first difficulty, we need to reduce the problem to a simpler problem of detecting these wavefronts instead of a full CSD wave. To address the second difficulty, we use higher-density EEG grids, and project the signal on the brain surface using a tool called “surface-Laplacian” [11]. Instead of deciding on the presence of CSD from the channels independently, we track wavefronts that move consistently across time with speed consistent with the range of CSD propagation speed. To obtain these wavefronts, we use displacement vectors which are known as “optical flows” and used in computer vision for object tracking [12] [13].

We test our algorithm on EEG data generated by simulating CSD propagation on the cortex using two models: (i) an ion diffusion-based “mesoscale” model of CSDs that builds on Tuckwell’s model of CSD propagation [14], with some modifications as described in Section II-A. Tuckwell’s model abstracts the physiological mechanisms of CSD propagation and helps us test our detection algorithm in an accepted model of CSDs. It makes a connection between the current understanding of CSD waves and our proposed algorithm. However, this model has a fixed width of suppression (~ 2 cm), and generates idealized ring shapes of CSD propagation (on planarized surfaces), while in practice, the width and the shape of CSD suppression can vary (by “width of CSD suppression” we mean the spatial width of the CSD wave parallel to the direction of propagation; see Fig. 4a). Therefore, we also use (ii) a more abstract model that suppresses regular brain activity (modeled as a random Gaussian process) to generate complex CSD patterns (e.g., a non-ideal ring shape, semi-planar wavefronts, and single-gyrus propagation). These patterns are simply imposed onto the random brain activity, and the generation is thus somewhat “artificial.” This artificial generation of CSD models allows us to tune suppression widths and shape of CSD waves, yielding a broader set of signals to test our algorithm on them. To generate EEG signals, the simulated CSD wave is projected to the scalp using subject-specific forward models (with 33,255 sources on the cerebral cortex (gray matter) and 40 and 340 sensors for low-density (LD) and high-density (HD) EEGs, respectively). We use MRI images from 4 subjects aged 18 to 74 years from the OASIS dataset^3^ to generate the forward models. Our results suggest that while low-density EEG grids can detect a significant fraction of CSDs, higher-density grids are required for detecting narrow CSD waves.

### A. Related works

In [10], Dreier *et al*. summarize the works of the Co-Operative Studies on Brain Injury Depolarizations (COSBID) group which utilize direct-current (DC) shift of ECoG signals to detect post-trauma CSDs. They show that in parts of the cortex without spontaneous neural activity (electrophysiologic penumbra), there is a specific type of spreading depolarization waves, called isoelectric spreading depolarization (ISD). ISDs can be recorded and visually observed using near-DC-ECoG signals, which contains the low frequency and near DC components of the recorded ECoG signals (below 0.5 Hz). This detection technique, however, could be hard to apply to noninvasive EEG signals where the preprocessing often filters out near-DC signals. We note that DC-coupled EEG recordings can be made from the scalp, and this can enable detection of CSDs using DC shifts in EEG [15], [16]. However, DC-coupled EEG is not commonly used today, thus we focus on high-frequency suppression in this work.

While ECoG signals are invasive, in [16], Hartings *et al*. show that CSDs can be detected noninvasively using EEG in patients with severe TBI. This work built on an earlier work of Drenckhahn *et al*. [15], who present similar results on patients with malignant hemispheric stroke and subarachnoid hemorrhage. Hartings *et al*. first identify CSDs in electrocorticography (ECoG) recordings through visual inspection, and also visually inspect time-aligned scalp EEG recordings to find amplitude depressions associated with the depolarization. They show that 81% of the identified events has manifestations in the “time compressed” (*i.e.*, downsampled) EEG recordings. The authors first downsample the EEG signals in time to visually observe electrical silences in EEG signals and then check if these silences spread spatially. However, in this work, the skull of the patients is fractured, potentially reducing the spatial blurring in the EEG signals. Further, the widths of CSD depressions are quite large in severe TBI, making these CSDs easier to detect noninvasively. Finally, the visual inspection technique used is not automated. All three limitations are overcome in our work.

A contrasting, and perhaps cautionary, recent work is that of Hofmeijer *et al*. [17], where the authors monitored 18 stroke and 18 TBI patients. Using 21 electrode EEG systems, no CSDs were observed through visual inspection. The authors speculate that, among author-notes reasons, this could be because of volume conduction and low resolution of the EEG system used. Because higher-density EEG systems continue to yield a higher spatial resolution [18], [19], a deeper understanding of what CSDs can be detected noninvasively through low and high-density EEG needs to be obtained experimentally. Our work provides a simulation-based understanding of this issue, and while an experimental validation is required, the simulations do suggest that low-density EEGs can miss narrow-width CSDs, consistent with the speculation in [17].

Towards developing automated algorithms, one related work is that of Gharibans *et al*. [20], who propose an automated algorithm to detect a slowly propagating gastric waves using noninvasive *electrogastrogram* (EGG). This detection algorithm, like ours, is based on consistency in speed and direction of propagation. However, our algorithm is more suited to the CSD detection problem because: (i) unlike in CSD propagation, there are no folds on the stomach surface to cause “disappearance” of the recorded wavefronts, and (ii) their algorithm utilizes spatial averaging over all electrodes, which limits the algorithm to detect time of spread, but not the spatial location. Similarly, in [21], Bastany *et al*. monitor spreading depolarization DC potential changes in rats using three different analyses, namely, spectrogram, bi-spectrogram, and pattern distribution. However, rat cortex also does not have any folds, and further, rats have thinner skulls than humans, causing less spatial blurring in EEG recordings.

### B. Paper organization

The rest of the paper is organized as follows. In Section II, we introduce our “mesoscale” model of CSD propagation that builds on a similar model of Tuckwell *et al*. [14], [22]. To complement Tuckwell’s model, we also introduce our models of complex CSD patterns that have varying CSD widths and patterns. In Section III we provide our automated CSD detection algorithm. In Section III-C, we quantify the performance of detection using low and high-density EEG on CSDs of varying widths of suppression, and for different head models. Finally, we conclude in Section IV, where we discuss limitations of the proposed algorithm and some possible directions for future work.

## II. CSD MODELING

In the following, we describe how we simulate the CSD wave on the cortex using a 2D “mesoscale” model, and perform forward modeling to project the signal onto the scalp to obtain simulated EEG data. To achieve realistic simulations of CSD waves, we use real brain models that have gyri and sulci. Fig. 2 summarizes the main steps of CSD modeling.

**Fig. 2:**
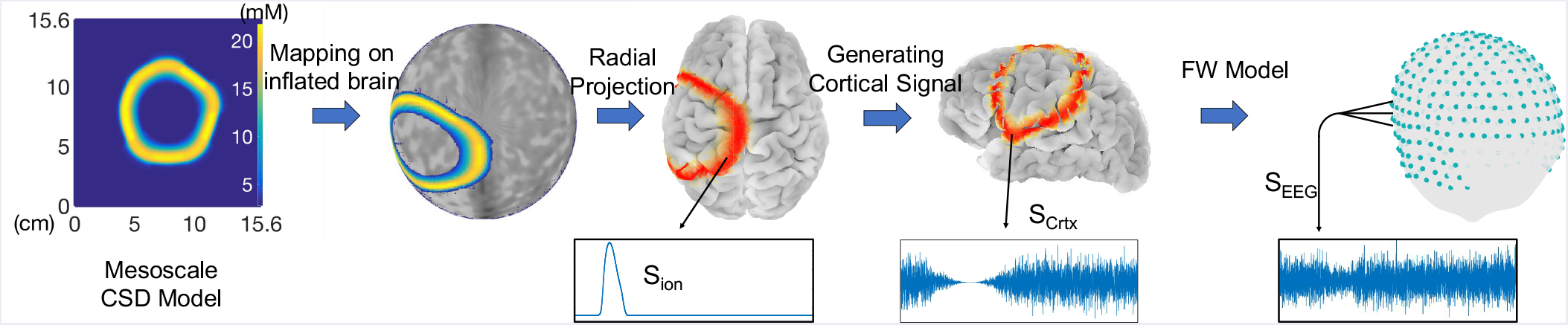
CSD Modeling: 4 main steps of simulating EEG signals obtained from CSD waves.

### A. 2D models of CSDs: a mesoscale model, and imposed complex CSD patterns

We borrow the 2D model of Tuckwell [22], which is based on extracellular space (ECS) and intracellular space (ICS) ion concentration changes. Tuckwell’s model is based on “reaction-diffusion” of ions and neurotransmitters. “Reaction” refers to the ion exchange between ICS and ECS, which is at the cell level and is the *microscopic* part of the model, and “diffusion” refers to the ionic propagation in the ECS and between neurons, which is the *macroscopic* part of the model. The resulting “mesoscale” model takes six components into account: 4 ions *K*^+^, *Ca*^++^, *Na*^+^, *Cl*^−^ and two neurotransmitters, one inhibitory which we call “*T_I_*,” such as GABA; and one excitatory (“*T_E_*“), such as glutamate. Among these six components, ICS concentrations of *K*^+^, *Na*^+^, and *Cl*^−^ are associated with the post-synaptic membrane, and ICS concentration of *Ca*^++^ and fluxes of *T_I_* and *T_E_* are associated with the pre-synaptic neuron. This mesoscale model consists of six coupled 2D parabolic partial differential equations (PDEs), which update the ECS concentrations of the 6 components on the 2D space and time as follows:

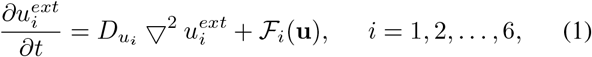

where **u**(*x, y,t*) is the vector of ECS and ICS concentrations, *D_ui_* is the diffusion coefficient of the corresponding ECS component, and 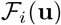 is the flux term of each component. To solve the PDEs in (1), we perform discretization using Euler’s method. Further details of this model are available in Appendices.

Fig. 5 shows a snapshot of the membrane potential (*V_M_*) in this mesoscale model at *t* = 15 min. During propagation of a CSD wave, *V_M_* gradually increases to a voltage which is greater than the threshold for neural firing (−55 mV). On the author-notes hand, CSD propagation is characterized by suppression of activity. This seeming inconsistency can be explained by noting that the neurons may not fire even as the membrane potential exceeds the threshold because they may not have enough energy. During CSD propagation, neurons lose a large amount of intracellular potassium, and consequently their electrochemical energy (stored in the ion concentration gradient) [5], which is needed to generate action potentials.

Some notable aspects of Tuckwell’s model are: (i) it does not account for cell swelling, which can increase the refractory period. However, the speed of CSD wave propagation is unaffected by cell swelling [23] even if the width could be affected. Thus, we also use author-notes CSD models where the width and the speed of depression are tunable parameters; (ii) the dynamics of calcium in the pre-synaptic region are related to the amount of transmitter release at the synaptic cleft (see Appendices), thus this model includes the dynamics of calcium [24]; (iii) the model instigates CSD using a local increase in the extracellular potassium concentration. In brain injuries, substantial intracellular potassium is released into the ECS of the injured part due to neural damage or death, which is believed to instigate CSDs. In fact, the amount of released *K*^+^ is proportional to the severity of the injury [25], [26]. Beyond injury, there may be author-notes factors as well that are involved in CSD instigation in neurological disorders.

Anauthor-notes assumption in [22] is the homogeneity of the medium in which the ions diffuse, which makes the shape of the generated CSD wave an idealized annular ring. Several factors can result in spatial heterogeneity of ion diffusions such as (i) the shape and the geometry of ECS that causes additional delay in ion propagation compared to a free medium, and (ii) presence of dead cells and fixed negative charges in the ECS, particularly in injured brains [27]. To make our simulations more realistic, we introduce heterogeneity in our model of the cortical medium, which is explained in detail in the Appendices. Fig. 3 shows the non-ideal ring shape of ECS concentration of potassium at three different time points during CSD propagation.

**Fig. 3:**
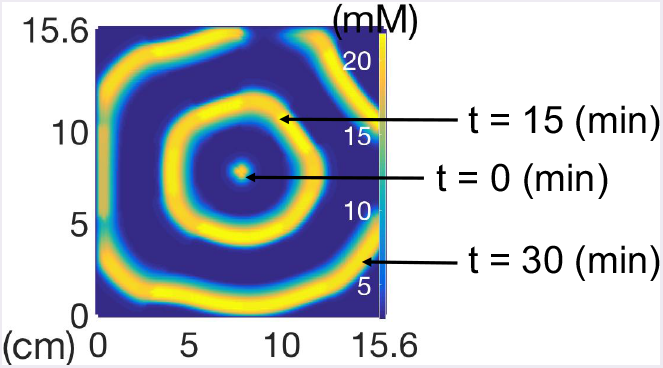
CSD propagation based on a heterogeneous mesoscale model. CSD waveforms are shown at *t* = 0, 15 and 30 (min).

In addition to the heterogeneous ring shape of propagation, CSD can have more complex patterns of propagation, namely, propagation on a single gyrus [28], [29], and spreading with multiple semi-planar wavefronts that split from an original wave [29]–[31]. Complex patterns of CSDs may appear when spreading depolarizations occur in energy compromised areas, e.g., lesions in brain [32]. To test the performance of our detection algorithm, we simulate these complex patterns of CSD on a 2D plane, similar to the 2D plane in the mesoscale model, and then project them on the cortex. Fig. 4a shows a CSD on left hemisphere with propagating semi-planar wavefronts with a width of suppression of 2.5 cm. Fig. 4b represents the propagation of a narrow CSD wavefront (5mm) on a single gyrus, which is highlighted in blue.

**Fig. 4:**
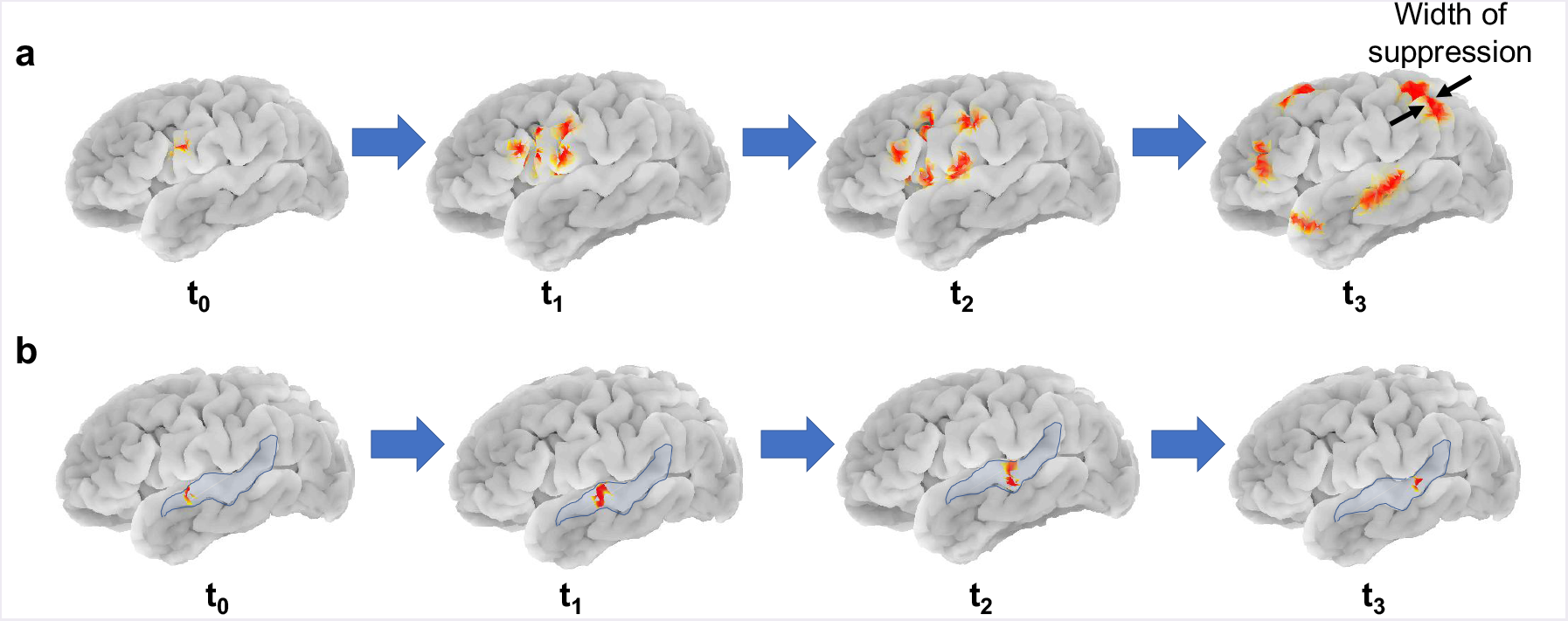
Complex patterns of CSD: a) Propagation of a CSD wave in the form of semi-planar wavefronts, shown in red, originating and splitting from an original wave with 2.5 cm width of suppression, b) Propagation of a narrow CSD wavefront (5mm suppression width), shown in red, on a single gyrus on the left hemisphere, highlighted in blue. These are complex patterns of CSD propagation suggested in [28]–[30]. In this work, we define the “width of CSD suppression” as the width of CSD wave parallel to the direction of propagation (see the last image in (a)).

**Fig. 5:**
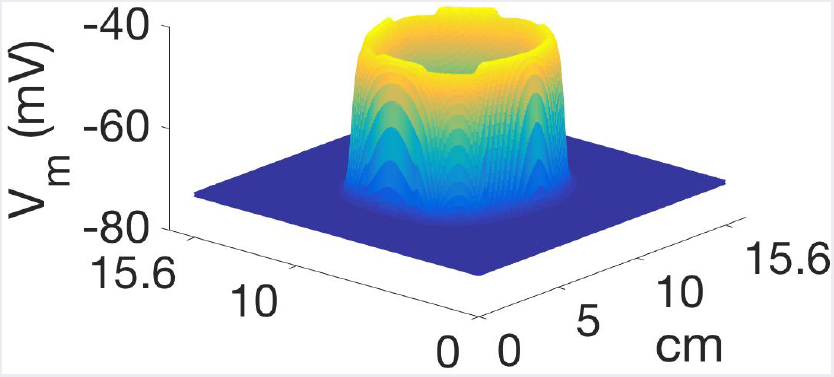
Membrane potential during CSD propagation at t = 15 min.

### B. Projection of cortical signals onto the scalp

To test the detection of CSDs using noninvasive scalp EEG, we first simulate a CSD wave on a real brain model and then obtain the resulting scalp EEG. To transition from our 2D model to real brain model, we perform mappings in four steps: (i) the 2D model is mapped onto a segment of an unrolled cylinder; (ii) the unrolled cylinder is projected onto a unit sphere, by radially contracting all points in a direction perpendicular to the axis of the cylinder (see Fig. 6)^4^; (iii) as an intermediate step, a unit sphere is used to represent the entire cortical surface through “inflation” of cortex (see Fig. 7). This intermediate inflated cortex is a smooth surface. Such inflation is readily performed using tools such as *Freesurfer*^5^ which is an open source software; (iv) the sphere (along with the CSD mapped onto it) is finally deflated to reassume the intricate gyrencephalic structure of the cortex. Fig. 7a shows a CSD generated on the left hemisphere. Since the 2D CSD from the mesoscale model is mapped onto an inflated spherical 3D model of the cortex, the CSD wave gets projected onto both sulci and gyri, not only on gyri. Hence, the spreading depolarization wave travels inside sulcal regions (see Fig. 8), and the time it takes for CSD to get out of a sulcus depends on the depth of the sulcus.

**Fig. 6:**
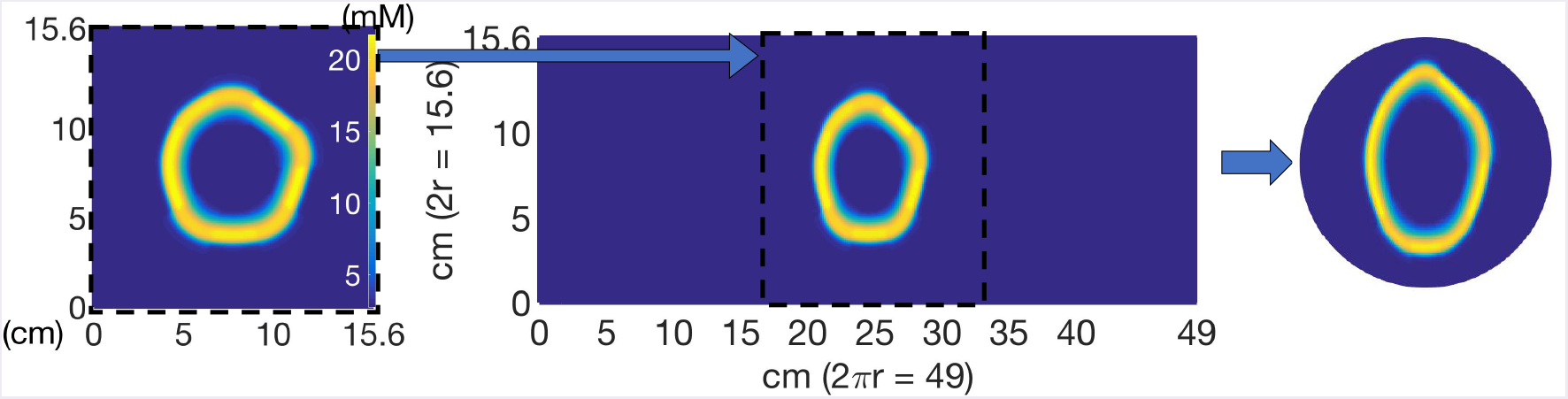
Mapping of the CSD wave onto the surface of a unit sphere (t = 15 min).

**Fig. 7:**
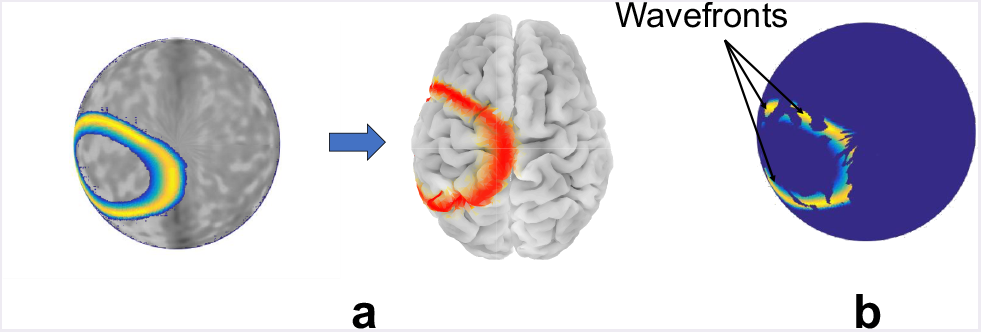
a) Mapping of the CSD wave from the inflated brain model onto the cerebral cortex of subject “OASIS2” (t = 15 min). b) “Electrical disappearance” of CSD waves inside sulci in the recorded cortical signals breaks the suppression pattern into multiple disconnected parts that we call “wavefronts.”

**Fig. 8:**
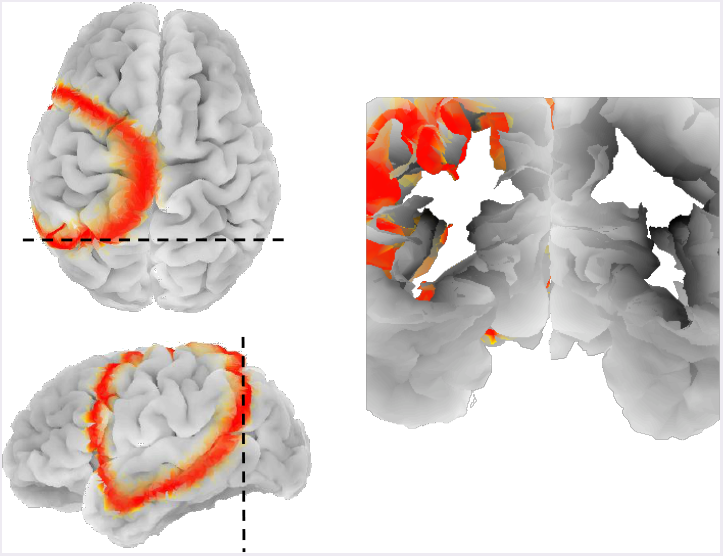
Coronal cross section of subject “OASIS2” showing how the CSD wave travels inside sulcal regions after mapping the 2D mesoscale model onto the inflated spherical model of the cortex.

To obtain the EEG recordings, we use a forward model based on the subject’s head model (obtained from MRI scans) to project the electrical activity of brain sources on to the scalp. In this paper, an open source MRI database (OASIS) is used to obtain real head models. We choose MRI image set of 4 healthy subjects in this dataset with different ages (OASIS1: 74, OASIS2: 55, OASIS4: 28 and OASIS5: 18 years old) and use *FreeSurfer* to process these MRI images and extract different layers of head, i.e. scalp, skull, cerebrospinal fluid (CSF) and brain (cerebral cortex). Fig. 9 represents the real head model and all of these layers for the subject “OASIS2”. Next, using a Matlab toolbox *FieldTrip* [34], which is designed for EEG data analysis, we generate a forward matrix with the first dimension equal to the number of EEG sensors on scalp and the second dimension equal to the number of electrical sources in brain (dipoles normal to the surface of cortex [11], [35]). In our simulations, we use 33,255 cortical sources and *N_ch_* (= 40 for low-density, or 340 for high-density) EEG electrodes on scalp. The EEG electrode locations are generated by projecting the spherical harmonic locations on the scalp and removing parts such as face and neck (Fig. 10).

**Fig. 9:**
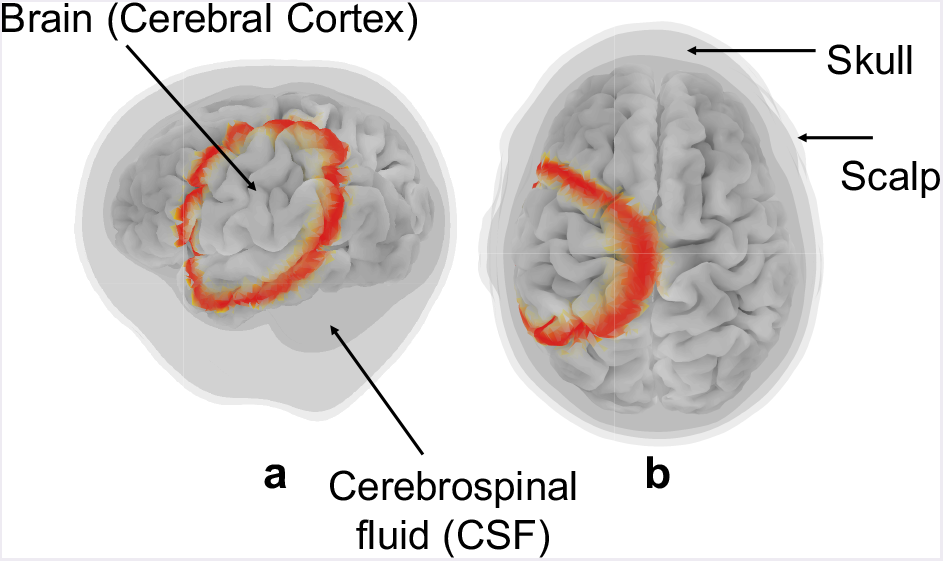
Real head model of subject “OASIS2”, including brain, CSF, skull and scalp layers which are extracted from its MRI image using open source *FreeSurfer* software: a) Left and b) top view of the head model.

**Fig. 10:**
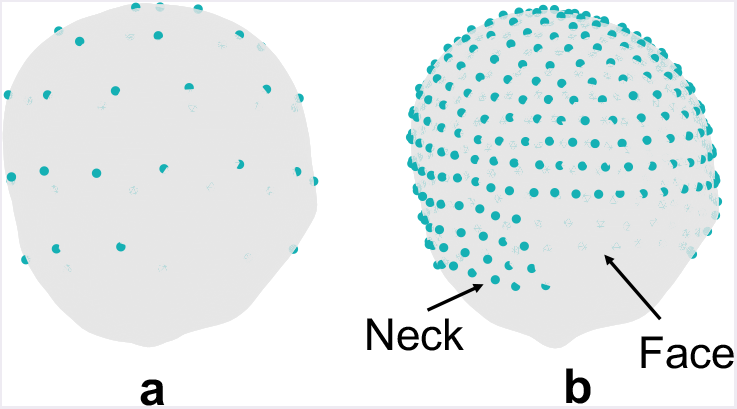
EEG electrode locations on a real head model (subject “OASIS2”) at spherical harmonic locations, face and neck electrodes are removed: a) *N_ch_* = 40 and b) *N_ch_* = 340 electrodes.

After constructing this forward model, the next step is generating electrical signals of CSD propagation on the cortex and project them on the scalp. To do so, we use two aspects of CSDs: i) During a CSD wave propagation, the activity of sources where the wave is passing is suppressed; and ii) Electrical activities of sources that lie within brain grooves (sulci) are hard to be recorded from the scalp, because of their depth, and also because some of these dipoles are parallel to the cortical surface. This breaks the CSD wave into disjoint parts that we call “wavefronts.” For the illustration purpose and to show how does the CSD wavefronts look like on the cortex we use the depth of sulci, extracted from MRI images, to create these breaks (See Fig. 7b). For the simulated EEG signals, the generated forward model automatically takes into account the “electrical disappearance” in sulci.

We simulate background electrical activity on the cerebral cortex during CSD propagation using normal random process and then suppress it at the locations of the CSD wave. We suppress the electrical activity using a smooth falling edge, and then gradually have it recover using a similar rising edge, with a variable delay^6^, after the CSD propagation. The delay is to model the fact that spreading depression usually outlasts spreading depolarization [10]. In this way, we utilize the spatiotemporal locations of CSD wavefronts in the “mesoscale” model to generate these suppressions in the electrical activity, and consequently, the main characteristics of CSD waves, including the speed of propagation and width of suppression, are reflected in the electrical signals. The simulated electrical signal used in our experiments lasts for an hour, and in the first 30 minutes, there is a CSD propagation that starts on the left hemisphere with the speed of 3 mm/min. Signal “*S_Crtx_*” in Fig. 11b is the simulated cortical signal. As the final step in this section, we obtain the EEG signals on the scalp (signal “*S_EEG_*”) by applying the subject-specific forward model to the simulated cortical signals. Fig. 12a shows the *S_Crtx_* signals at four different source locations on the cortex with the temporal locations of CSD suppression are marked by arrows, and Fig. 12b represents the resulting *S_EEG_* signals with corresponding suppression locations. Visual comparison of our simulated signals with the actual ECoG and EEG recordings of CSD propagation (e.g., [16]) suggests that our simulated electrical signals are a reasonable approximation of CSD waves.

**Fig. 11:**
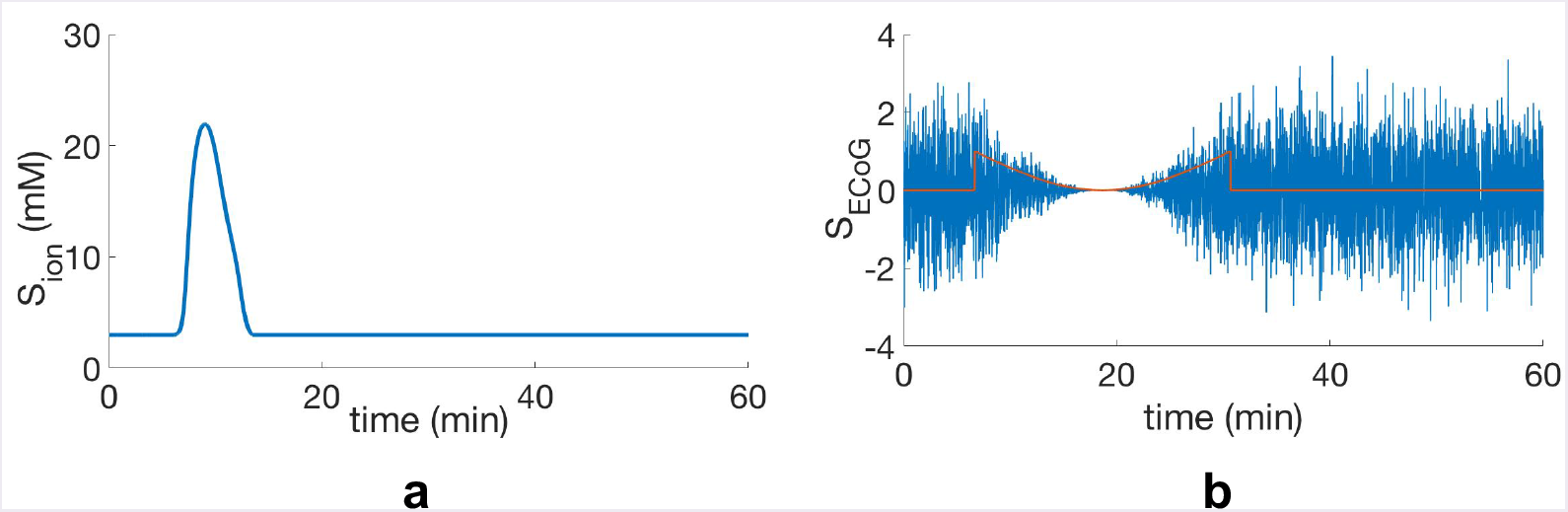
Generating electrical signals of CSD waves on the cortex: a) Extracellular potassium concentration during CSD propagation at one of the sensor locations (*S_ion_*), b) Background electrical activity (*S_Crtx_*), that we generate using a normal random process (*σ* = 1) and then suppress at locations of the potassium ion concentration peak in (a) by means of a smooth suppression pattern.

**Fig. 12:**
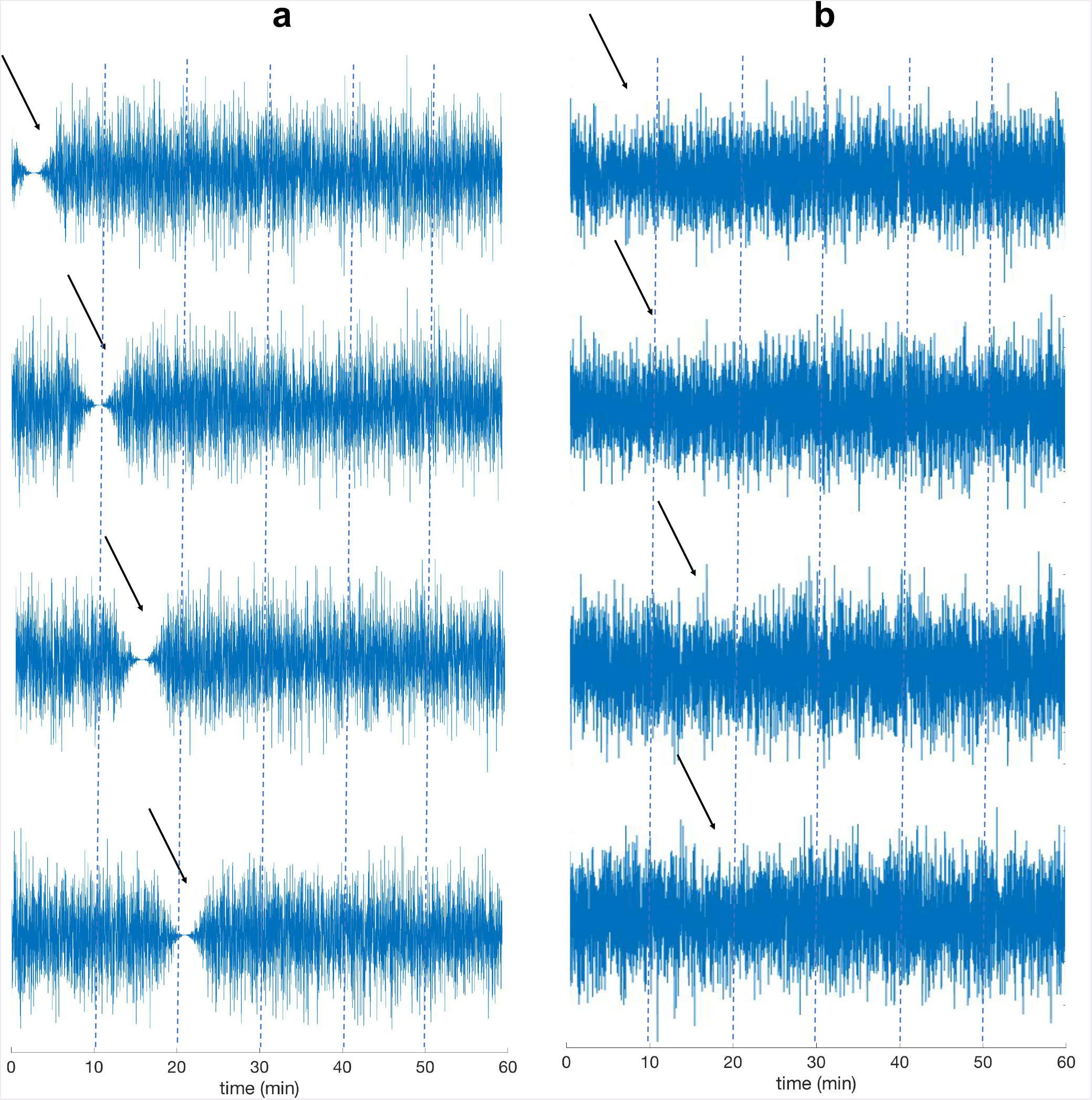
The simulated *S_Crtx_* and *S_EEG_* signals. The signals of this figure contain CSD suppression at *t* = 5,10,15 and 20 min. In the *S_Crtx_* signals shown in (a) the suppressions can be visually detected, while in (b) because of the attenuation of high spatial frequencies in *S_EEG_* the CSD suppressions hardly can be identified.

## III. DETECTION ALGORITHM

Detection of CSDs from EEG recordings is challenging because of the following reasons: (i) as explained in the previous section, EEG is less sensitive to activity inside a sulcus, breaking the recorded wave into multiple wavefronts; (ii) the decay of high spatial frequencies as the signal passes through skull and soft tissue. This makes the detection of CSDs with narrow width and/or small overall area of suppression hard, e.g., detection of CSDs with semi-planar wavefronts, which may affect only a single gyrus, is difficult using EEG; (iii) uncertainty in the point of initiation of CSD makes the detection challenging; and (iv) different speeds of propagation in different brain regions due to the spatial heterogeneity of the diffusivity of ions involved in the CSD propagation (see the “heterogeneous model” in Appendices).

To overcome these challenges, the key idea is to identify and track CSD wavefronts and stitch together data across time and space to detect the CSD wave. In doing so, we first pre-process the simulated EEG signals to obtain a sequence of interpolated 2D spatial maps, from which we extract spatiotemporal coordinates of suppressed activity. Because a CSD wave manifests as traveling wave of suppressed brain activity on the cortex, we track suppressions that spread across time and space and score the wavefronts based on their orientations and speeds to classify the propagating waves and detect CSDs. In the following sections, we explain each of these steps in detail.

### A. EEG pre-processing

The goal of this step is to extract the neural suppression pattern out of the simulated EEG signals, as these signals may have a low spatial resolution. Also, we partially compensate (using Laplacian spatial filtering) for the attenuation of high spatial frequencies caused by layers in between the electrical sources of the cortex and the EEG electrodes on the scalp surface, such as cerebrospinal fluid (CSF), skull and scalp.

Pre-processing the EEG signals helps improve the signal to noise ratio (SNR) of CSD suppressions and obtain a sequence of 2D spatial maps, on which we later apply image processing techniques for CSD detection. All of the following preprocessing steps are summarized in Fig. 13.

**Fig. 13:**
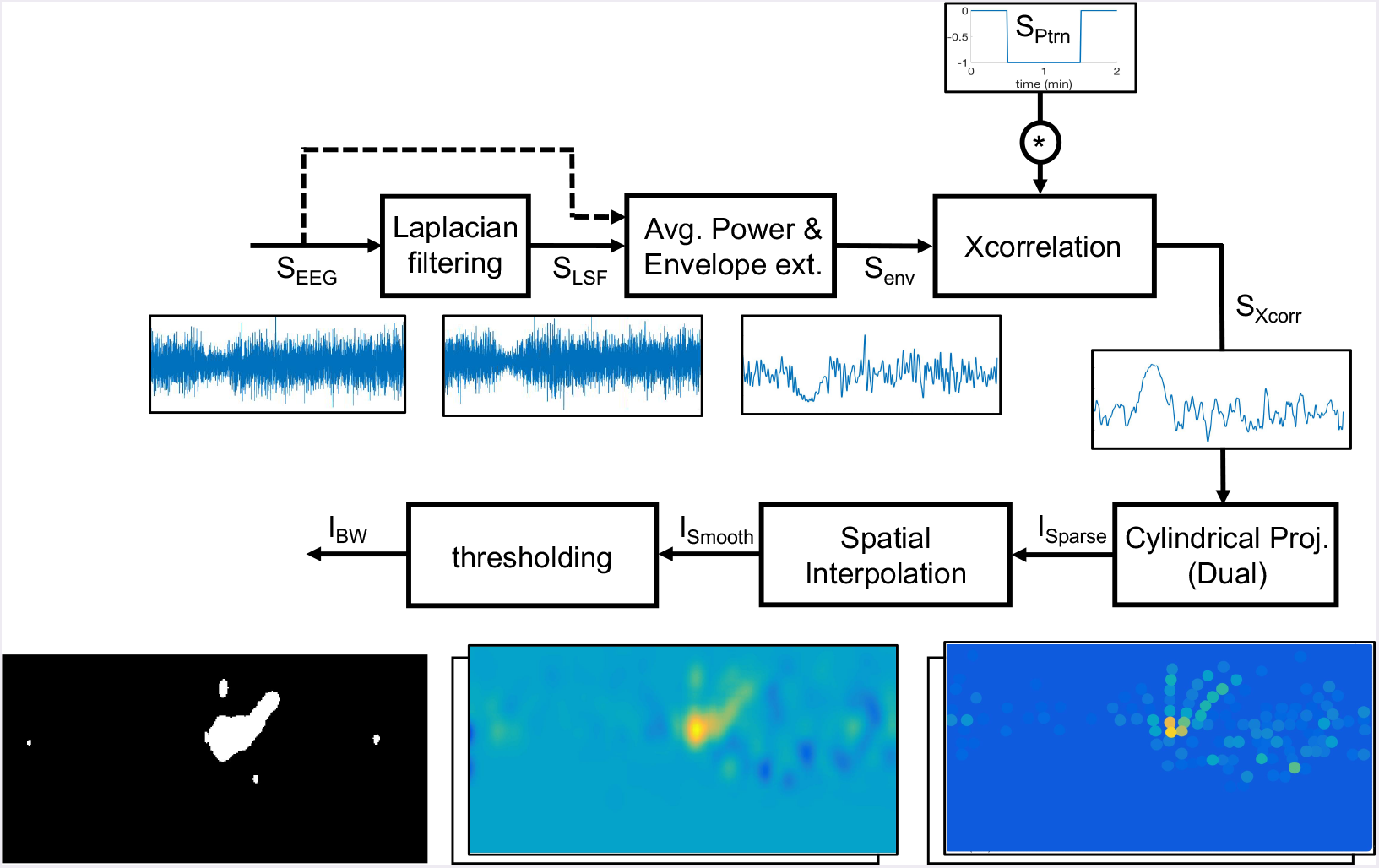
CSD pre-processing: main steps of extracting spatial maps of CSD suppressions out of simulated signals on scalp (*S_EEG_*).

*Laplacian spatial filtering:* Laplacian filter is an approximation of the 2D spatial second derivative that is a good approximation for high-density EEG. Operationally, it extracts localized activity with high spatial frequencies [36] [37]. Since narrow CSD waves have high spatial frequencies, Laplacian filter is a good choice to extract the suppressions from the EEG signals and overcome the attenuation of high spatial frequencies in these signals. In contrast, for wide CSDs Laplacian filtering may not help. To not make any assumptions on the width of CSD in the detection process, we apply our detection algorithm on signals both before (“*S_EEG_*”) and after (“*S_LSF_*”) applying the Laplacian filter, and make a decision based on both. As the electrode distance increases, the Laplacian filter becomes less sensitive to the highly localized activity, e.g., narrow CSDs [36]. Therefore, we only apply the filter to high-density EEG electrodes (*N_ch_* = 340). To apply the Laplacian filter on the *S_EEG_* signal at each electrode location, we consider its neighboring electrodes in radial distance of 3cm and update the voltages of electrodes as follows [36]:

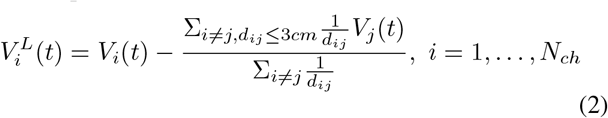

where *V*_i_ (*t*) and 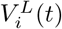 are the voltages of signal *S_EEG_* and its Laplacian (*S_LSF_*) respectively at electrode *i* and at time *t*, and *d_ij_* is the radial distance between electrodes *i* and *j*. Fig. 14 shows how the use of Laplacian helps in extracting a narrow CSD suppression from the signal *S_EEG_*.

**Fig. 14:**
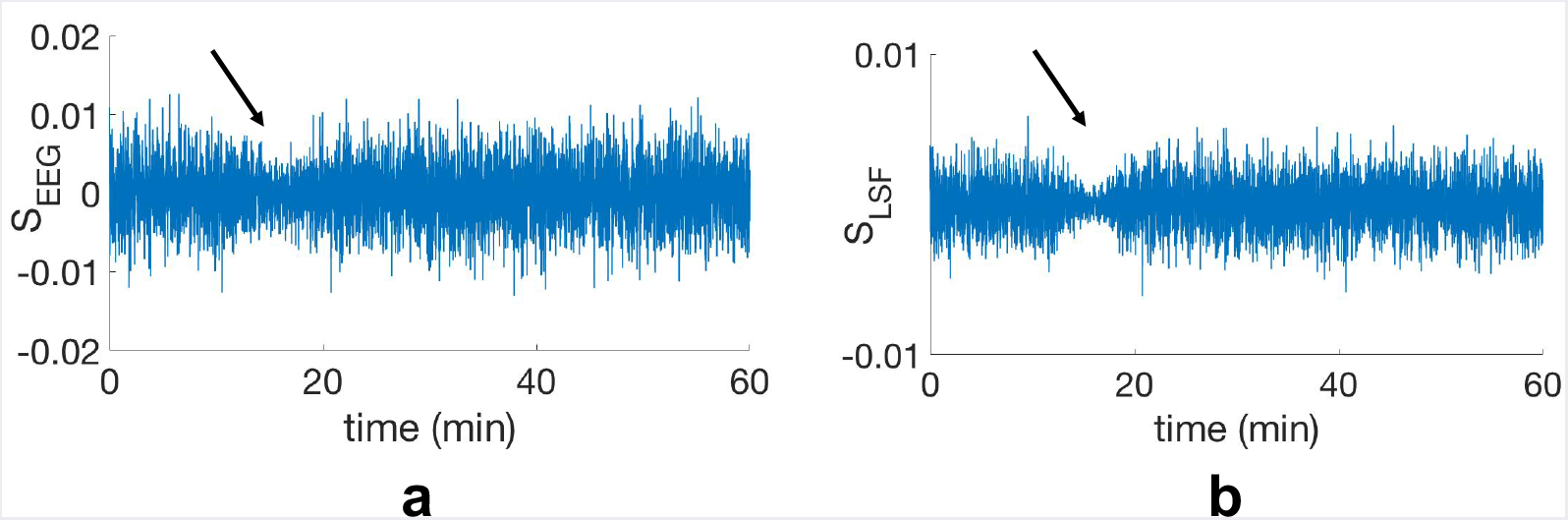
Weighted spatial Laplacian filtering of EEG signals to extract narrow suppression patterns of CSD: (a) shows an example of simulated EEG signal before (“*S_EEG_*”) and (b) shows after applying the Laplacian filter (“*S_LSF_*”). In this example, the width of CSD is 2 cm.

*Envelope extraction and cross correlation:* In this step, we calculate the average power of EEG signals using a sliding time window for all *N_ch_* channels. We arbitrarily assign a width of 14 seconds (20 samples) to this time window, which is small compared to the temporal width of CSD suppression. This low pass filtering temporally smoothens the EEG power. Signal “*S_PW_*” in Fig. 15a shows this average power.

**Fig. 15:**
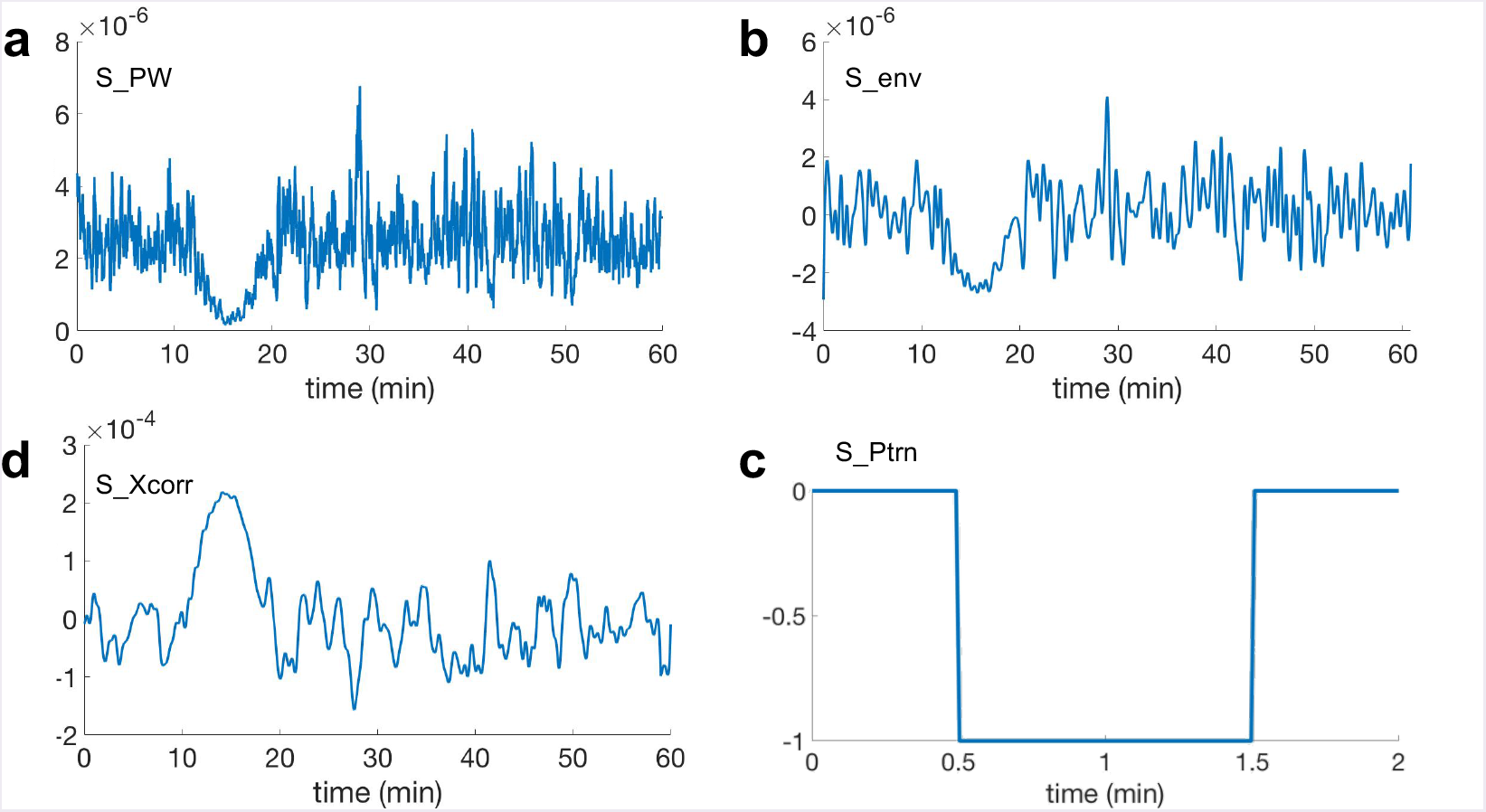
Preprocessing of EEG signal: a) *S_PW_* : smoothened EEG power using an averaging time window of 14 seconds, b) *S_env_*: zero-biased peak envelope extracted signal from SP W with minimum peak distance of 14 seconds, c) *S_Ptrn_*: “negative pulse” pattern with temporal width of 1 minute, d) *S_Xcorr_*: cross correlated *S_env_* with the *S_Ptrn_* signal. The amplified peak in (d) indicates the presence of suppressed brain activity at *t* =15 min.

Next, we perform envelope extraction. We use peak envelope extraction method, proposed in [38] and implemented in [39], which detects and extracts local maxima ^7^. In Fig. 15b, signal “*S_env_*” is the extracted envelope with zero DC offset. It is necessary to remove the DC offset of signal *S_env_* for the next step, which is the amplification of the CSD suppression. We do this amplification using the cross correlation of signal *S_env_* with a pattern of a negative step function (signal “*S_Ptrn_*” in Fig. 15c). We exploit the fact that CSD suppression causes a reduction in the power of EEG signals. This power reduction forces the signal *S_env_*, which has zero DC offset, to go to the negative values. We choose a 1-minute wide negative pulse for *S_Ptrn_* in a way to be able to extract all types of CSDs which may have from 1 to 64 minutes long suppressions at any location [4], [16], [40]. The output signal after the envelope extraction and amplification of the silencing is shown in Fig. 15d as signal “*S_Xcorr_*”.

*Cylindrical projection:* Our detection algorithm utilizes *optical flow*, a video-processing technique designed for 2D images. In order to obtain 2D spatial maps of pre-processed EEG data (*S_Xcorr_*), we project the scalp EEG electrode locations onto a 2D plane using cylindrical projection and assign the signal *S_Xcorr_* to each corresponding electrode (image *I_Sparse_* in Fig. 16).

**Fig. 16:**
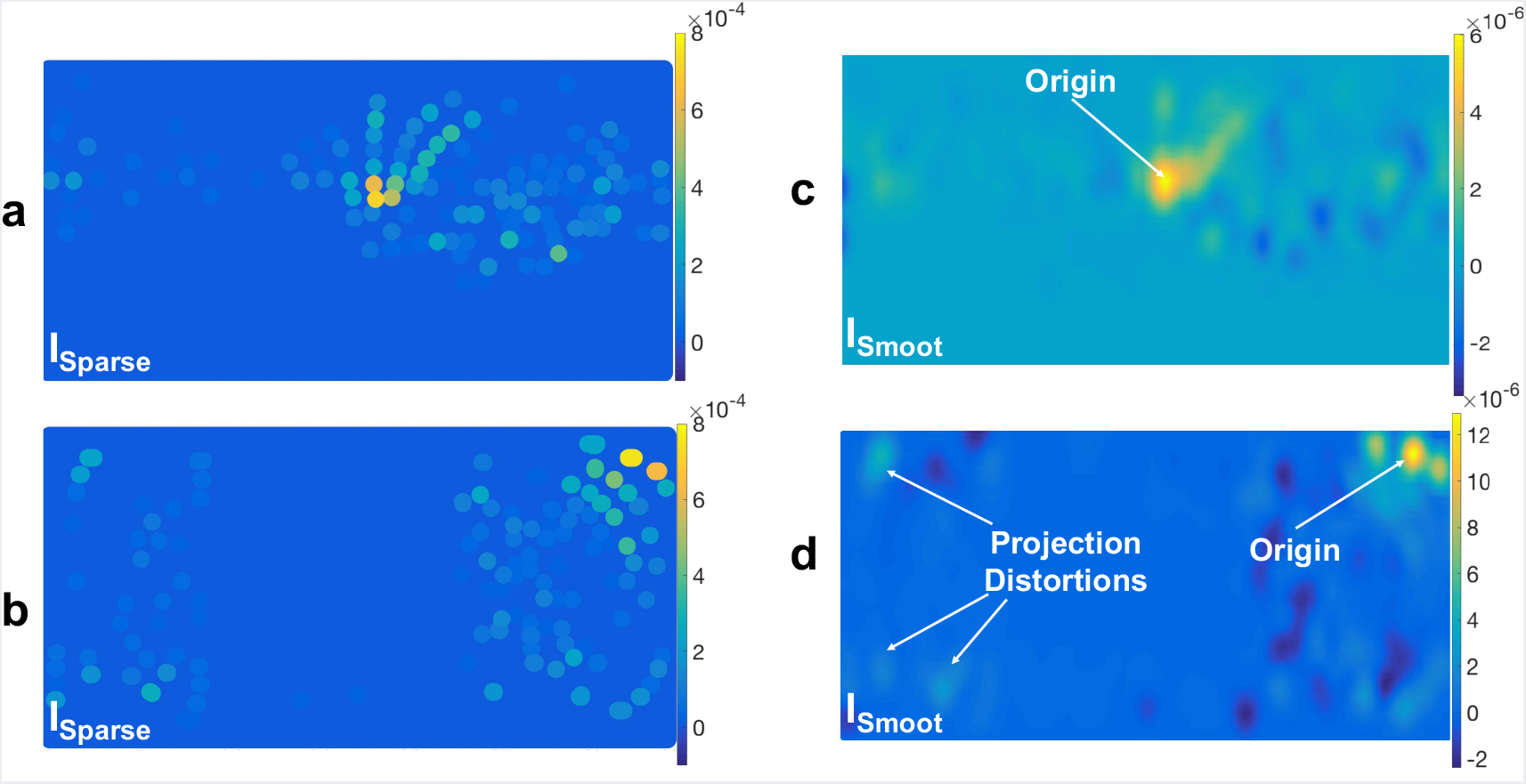
Cylindrical projection of preprocessed scalp EEG signals (*S_Corr_*) onto a 2D plane (*I_Sparse_* in (a) and (b)) and their spatial interpolation (*I_Smooth_* in (c) and (d)) using a Gaussian spatial kernel with *σ* = 6.8 mm. In *I_Sparse_* each dot indicates an EEG electrode location and its color indicates the corresponding amplitude of *S_Xcorr_* (t = 10 min): a) a cylindrical projection which is oriented in a direction that captures the CSD wave in the middle of the plane and (c) is its spatially interpolated image; b) a cylindrical projection which is oriented normal to the previous one and, in this example, it captures the CSD wave near the boundary of the plane and causes distortion in the wavefronts and (d) is its corresponding interpolated *I_Smooth_*.

To address the uncertainty in the location of the focal point of CSD waves, we take two different cylindrical projections which are oriented normal to each author-notes. Fig. 16 illustrates an example of this dual cylindrical projection. By using these dual projections, we ensure that the CSD wave is represented without distortion at the edges of the 2D plane. In Fig. 16b, the starting point is near the upper right corner, and the shape of the CSD wave is quite distorted, which makes the tracking of wavefronts unreliable. In Fig. 16a, this problem is resolved, and the wave appears near the center of the 2D plane. Therefore, we use both cylindrical projections in our detection algorithm, but keep track of CSD propagation based on the one that captures the CSD wavefronts (if any is present).

*Interpolation and thresholding:* The 2D spatial map of the scalp simulated signals that we obtain using the previous steps (*I_Sparse_* in Fig. 16a and b) is spatially sparse due to the limited number of EEG electrodes, even more so with low-density EEG. Therefore, we apply a 2D Gaussian kernel with *σ* = 6.8 mm to spatially interpolate and create smooth images out of these sparse 2D maps. Image *I_Smooth_* in Fig. 16c and d shows these interpolated 2D spatial maps.

Having these 2D spatial maps, we apply a two-step thresholding process to obtain binary images: a global and a local threshold. We first apply a global threshold to reject pixels with minimal values in *I_Smooth_* . Concretely, we assign zero to pixels with values lower than 40% of the global maximum (the average of highest pixel values among all images). As a second thresholding step, for each smoothened image, we set the pixels for regions whose magnitudes lies within upper 60% of pixel value range in that image (local threshold). This threshold is kept not too high to reject valid CSD wavefronts, even allowing for a relatively high “false-alarm” rate where positive detections will be rejected later through stitching data across time and space. Fig. 17 shows the resulting binary image (*I_BW_*). The regions of higher magnitude correspond to regions of suppressed electrical activity, and they appear as connected components in *I_BW_*. Some of these connected components are parts of the CSD propagating wave, and author-notess correspond to non-CSD evanescent waves, which we expect to reject in the ensuing processing.

**Fig. 17:**
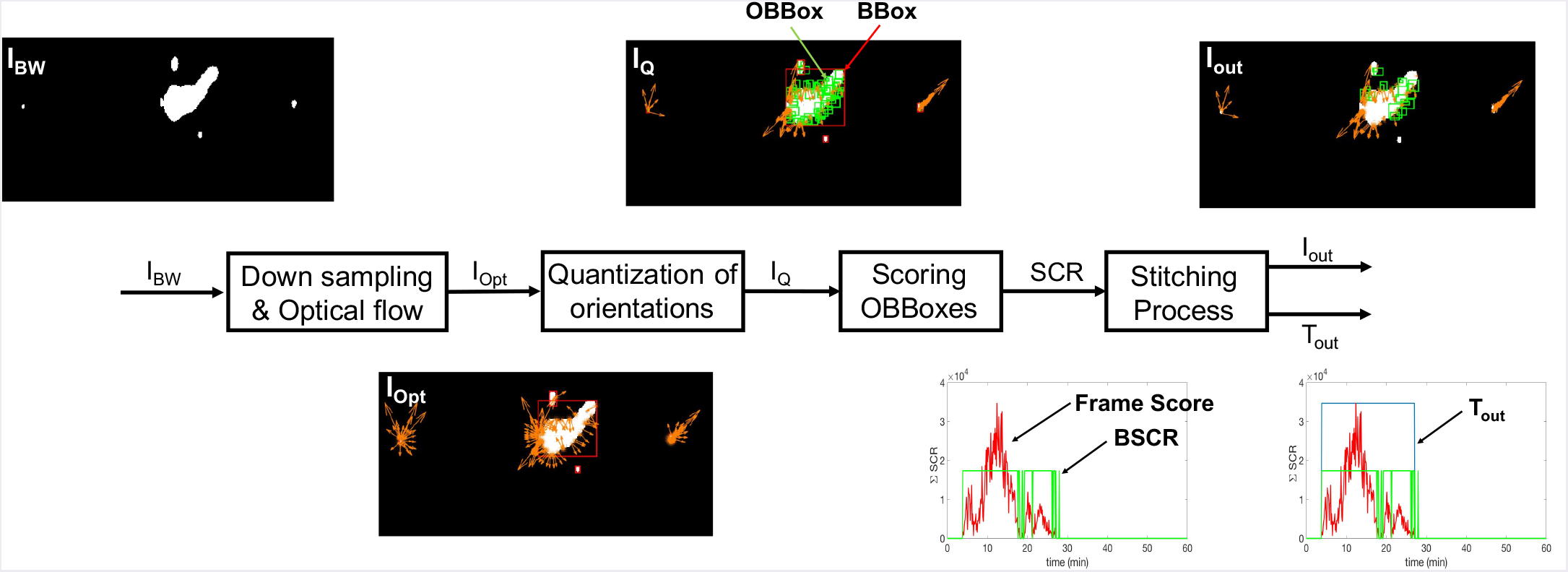
The steps of our CSD detection algorithm.

### B. Tracking of wavefronts and stitching data across space and time for an overall detection decision

In order to identify whether the wavefronts belong to a CSD propagation, we track the motion of wavefronts across frames using optical flows of pixels, followed by computing a score function based on the direction of propagation and speed and through a stitching process across time and space. Thresholding on the overall score function is used to detect the CSD waves. Fig. 17 shows different steps of the detection algorithm and the connection between them.

*Downsampling and calculating optical flow:* Optical flow is a technique used in computer vision to calculate the movement of objects between frames based on the spatiotemporal brightness variations and is used in many tasks such as motion estimation, object tracking and segmentation [12], [13]. We leverage optical flow to obtain the speed and orientation of the wavefronts’ motions. We use the Horn-Schunck algorithm for computing optical flows, which is proposed in [12] and implemented in [41]. However, since the speed of CSD propagation is very slow, we temporally downsample the binary images by means of sub-sampling every ten frames (every 7 seconds), to make it easy for the optical flow algorithm to capture the CSD wavefronts’ movement across frames.

Image *I_Opt_* in Fig. 19c shows the motion vectors as a collection of arrows where the length of each arrow corresponds to the magnitude of velocity and its orientation gives the direction of propagation of the corresponding pixel in *I_BW_*.

*Quantization of orientations:* As shown in Fig. 19c, in each binary image there are some connected components. In this step, we assign a bounding box to each connected component, and we call it “BBox” (red boxes in *I_Opt_*). To reduce the computational cost and focus on the larger connected components, we reject the bounding boxes with an area less than 20 *mm*^2^. Based on the size of the Gaussian kernel that we use for spatial interpolation (*σ* = 6.8 mm, see Section III-A), this is a reasonable threshold for the size of bounding boxes.

Next, we compute a histogram of the orientation of optical flows for each of these bounding boxes. Fig. 18b shows an example of this histogram. The histogram contains 12 bins of 30° each. Based on this histogram, the prominent directions of propagations can be extracted, e.g., if there are two significant directions of propagation in a bounding box, it indicates that the wavefront might split into two parts during the next few frames. To extract these prominent orientations, we apply a thresholding operation on each of these histograms at 50% of the largest population among the bins. For the remaining optical flows, we quantize the orientations based on the central point of their corresponding bins in the histogram, e.g., if the orientation lies in the interval of 60° to 90°, its new value becomes 75°. Fig. 18c shows these quantized optical flows.

**Fig. 18:**
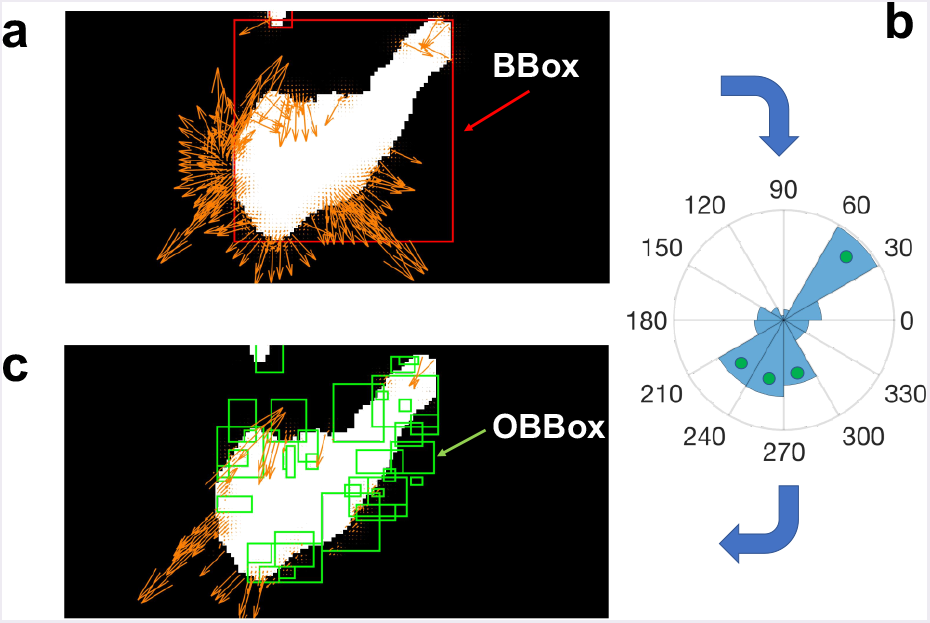
Quantization of optical flows based on a histogram of orientations: a) a BBox and its corresponding optical flows (*I_Opt_*); b) the calculated histogram of orientations for optical flows in *I_Opt_*. Green circles indicate the selected bins after applying a threshold at 50% of the maximum population; c) Quantized optical flows based on the selected bins in (b). Bounding boxes of the quantized orientations (OBBox) are shown in green.

*Orientations bounding boxes (OBBox):* In this step, we define a new set of bounding boxes based on the quantized orientations. For each value of orientation, we find the optical flows that are connected through pixels and have similar orientations, and we assign a bounding box to each of them (green boxes in Fig. 18c). We name these new bounding boxes “OBBox”.

We apply a threshold based on the size of these OBBoxes, similar to the threshold for BBoxes as we explained in the previous section, to reject very small OBBoxes and focus on the significant parts of the wavefronts which move together in the same direction.

*Scoring OBBoxes based on the consistency of propagation:* In this step, we score these OBBoxes based on the consistency in the direction and speed of propagation. Since each OBBox contains optical flows with similar orientation, we consider this orientation as the direction of propagation of OBBox, and we assign the average magnitude of the optical flows inside each OBBox as its speed of propagation. Our scoring algorithm consists of two main parts: (i) define the spatiotemporal neighbors for each OBBox; and (ii) assign a weighted score to each OBBox based on direction and speed of propagation.

First, for each OBBox we define its neighbors as the OBBoxes which lie within a radius of 2 cm (center-to-center distance) and temporal range of about 2 minutes (20 frames before and 20 frames after the current frame). Since a CSD wave propagates slowly (8 mm/min at the maximum [16]), inside this window of 2 minutes, we can expect the CSD wavefronts to have a displacement of less than 2 cm. This explains the reason for choosing the values above as the spatiotemporal range of neighbors.

As the second step in the scoring algorithm, we check if the speed of each OBBox is consistent with that of a CSD wave (1-8 mm/min). If so, we search its spatiotemporal neighbors, as defined in the first step, to find matching bounding boxes, i.e., an OBBox with the same direction of propagation and an average speed of 1-8 mm/min. We define the score of each OBBox (“SCR”) as the sum of the area of its matching neighbors. Thus, we now have a weighted score function for OBBoxes. Note that although the cylindrical projections (Section III-A) reduces the speed of propagation for the projected wavefronts in some parts of the 2D plane (depending on the orientation of the cylindrical projection), the upper bound of 8 mm/min on the speed of CSD still holds and there is no faster CSD propagating wavefront on the 2D spatial maps after projection.

In addition to the spatiotemporal scoring (SCR), we check the temporal consistency of this score function, i.e., whether these matching OBBoxes are well distributed over the temporal range of 20 frames, or just concentrated in a few frames. This helps us differentiate and reject the short-time non-CSD evanescent waves from the consistent and slowly propagating CSD wavefronts. We assign a binary score to each of the 40 neighboring frames; if a frame contains one or more matching OBBoxes, we assign 1 to this score; author-noteswise, we assign a 0. We call this the “temporal score”. If less than 60% of these frames have non-zero temporal scores, we reset (make zero) the spatiotemporal score of corresponding OBBox. This scoring algorithm allows us to extract the OBBoxes that are candidate pieces of a CSD wave. However, there is still room to improve the results regarding rejecting false alarms, using exploiting the temporal characteristic of CSD propagation, i.e., the duration of propagation. This is done in the next step.

*Stitching process and the final decision on detection:* In this step, we make the final decision based on the spatiotemporal scores of OBBoxes. First, to reject the boxes with minimal scores, we apply a spatial threshold, i.e., the 10% of the maximum available score at each frame. Also, we reject the frames (and all of the **OBB**oxes within them) with a total score of less than 10% of average scores among all frames. Here, the frame score is simply the sum of the scores of all OBBoxes within it. Fig. 19b shows this frame score (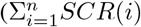, where n is the number of total frames) after thresholding.

**Fig. 19:**
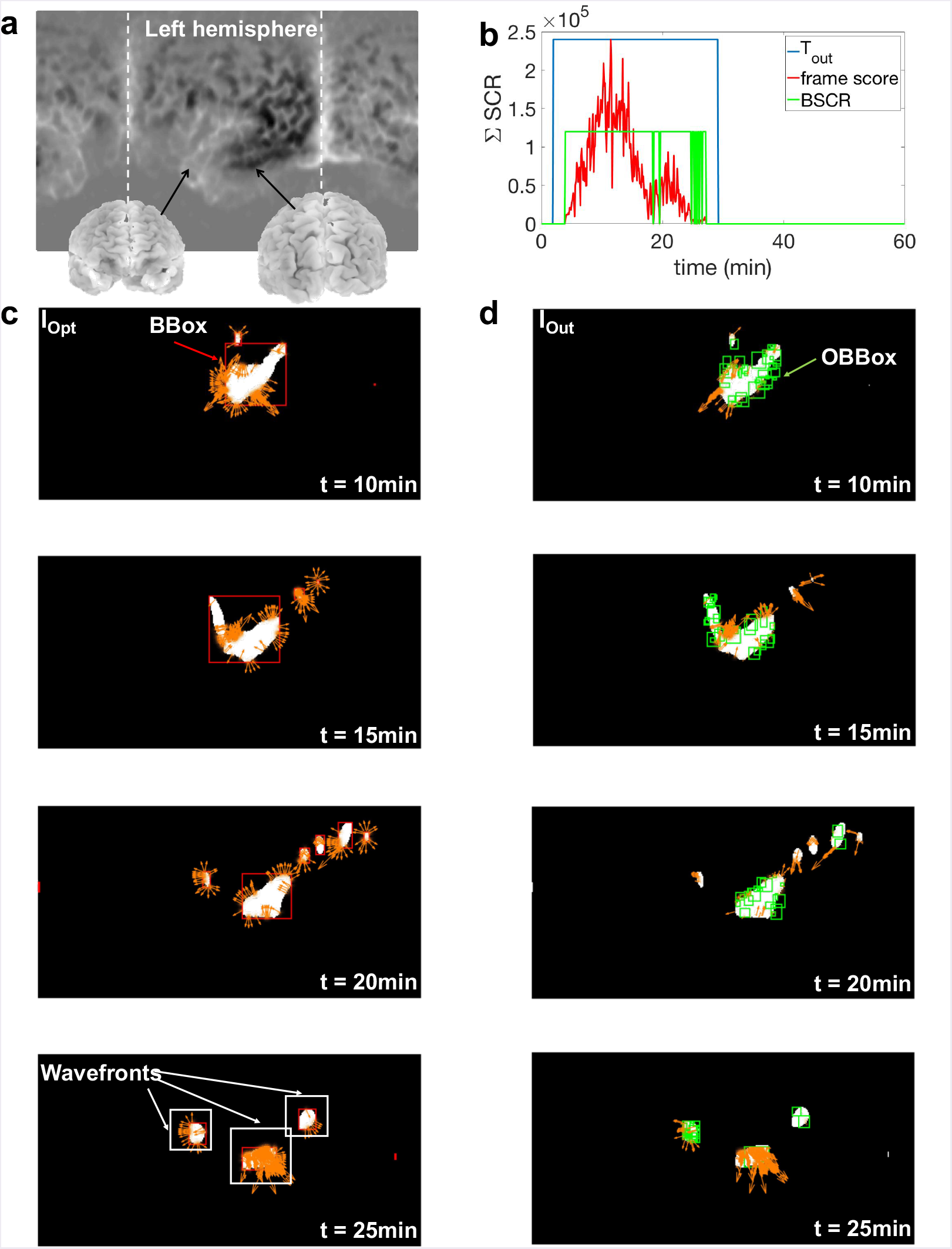
a) Locations of different parts of brain for the binary images *I_Opt_* and *I_out_* in this example. In this figure, the left hemisphere is located in the middle where the CSD starts to propagate; b) temporal output of the detection algorithm (*T_out_*) for the simulated CSD using the mesoscale model (2 cm wide); c) *I_Opt_* which shows optical vectors of CSD “wavefronts” in addition to the BBoxes of connected components (red boxes); d) *I_out_* in which the detected CSD wavefronts are marked using multiple green bounding boxes (OBBox). Based on the consistency in the direction and speed of the wavefronts, these OBBoxes are scored and selected.

We stitch the frames as follows: we slide a time window of 10 minutes over the frames and calculate the number of frames with non-zero scores (“*BSCR*”=1, see Fig. 19b) which lie inside this time interval. If this value is greater than half of the total number of frames inside this time interval (5 min), we select the middle frame. Fig. 19b shows these selected frames as a group of connected time indices using a binary function “*T_out_*” (1=selected frame). We check the temporal length of these connected frames to be more than 5 minutes. After this step, we keep the OBBoxes of the selected frames and reject author-notess. The final outputs of this detection algorithm are: i) the spatial location of the remaining OBBoxes, which indicates the location of CSD wavefronts in each frame (*I_out_* in Fig. 19d); and ii) the temporal locations of these selected frames as the time indices of CSD propagation in the simulated dataset (*T_out_* in Fig. 19b). The output of our detection algorithm using HD-EEG simulated signals (*N_ch_* = 340) on a wave generated using Tuckwell’s model (Section II-A) is shown in Fig. 19.

### C. Detection results: comparing low and high-density EEG

We apply the detection algorithm on four different real head models, which we extract from the OASIS dataset of MRI scans. CSD suppression widths for TBI can range from 0.8 to 6.4 cm [4], [16], [40], and the range could be broadened by including migraines (very short suppressions). Thus, we test the performance of our algorithm for a wider range of widths of suppression, namely 0.3 to 6.4 cm. Here we explore how high-density EEG could improve the performance of CSD detection concerning the minimum detectable width of suppression.

We apply the detection algorithm on simulated signals from both low-density (LD) and high-density (HD) EEG grids (see Fig. 10). For the LD-EEG, we use 40 electrodes (*N_ch_* = 40), and for the HD-EEG, we use the same 340 electrodes (*N_ch_* = 340) as used in the previous sections. For both scenarios, we start with a CSD wave with a large width of suppression and reduce it to find the minimum detectable width of CSD. However, the mesoscale model of CSD, which is proposed in section II, has a fixed width of CSD and is not appropriate for this experiment. To address this issue we “artificially” (i.e., without using Tuckwell’s model) generate CSDs on the real brain using the simulation of a propagating non-ideal ring-shaped wave emanating from a focal point.

Following the steps of CSD modeling in Fig.2, we first generate CSD waves on a 2D plane. In this 2D image, there are seven sectors of 51.4° each to which we assign different speeds of propagation in the range of CSD propagation’s speed (1-8 mm/min). In this simulation, we use the head models of 4 subjects and based on section II-B, we generate two forward models for each subject, one for the case of LD-EEG simulations (40 sensors and 33,255 sources) and the author-notes for the HD-EEGs (with 340 sensors and 33,255 sources). Following steps in Section III, we find the minimum widths of CSD waves that are still detectable by our algorithm. When using HD vs. LD-EEG, there are two differences to note: (i) we only apply the Laplacian filter to HD-EEG electrodes because the Laplacian is only effective at high densities [36]. HD-EEG captures cortical signals at a higher spatial resolution relative to LD-EEG [18], [19], and Laplacian filter can extract a high-spatial-frequency view of cortical activity; and (ii) because the spatial resolution of LD-EEG is low, we use larger Gaussian kernel (*σ* =13.8 mm) for spatial interpolation to obtain smooth 2D maps (*I_Smooth_*).

**TABLE I:** Minimum detectable widths of CSD using low-density (LD) and high-density (HD) EEG electrodes.

| Subject | EEG | Interpolation ($\sigma$ mm) | Minimum detectable width of CSD (cm) |
| --- | --- | --- | --- |
| OASIS1 | HD | 6.8 | 0.4 |
|  | LD | 13.8 | 1.0 |
| OASIS2 | HD | 6.8 | 0.3 |
|  | LD | 13.8 | 1.1 |
| OASIS4 | HD | 6.8 | 0.4 |
|  | LD | 13.8 | 1.4 |
| OASIS5 | HD | 6.8 | 0.6 |
|  | LD | 13.8 | 0.9 |
| Avg. | HD | - | $0.43 \pm 0.13$ |
| | LD | - | $1.1 \pm 0.22$ |

The results of these simulations are provided in Table I. The average minimum detectable CSD-width using LD-EEG is 1.1 ± 0.22 cm, which covers the majority, but not all known widths of CSDs. However, the average minimum

detectable CSD-width using HD-EEG electrodes is 0.43 ± 0.13 cm, which covers almost all types of CSD waves. Fig. 20 shows the output of CSD detection (*T_out_*) for narrow (1 cm) and wide (6.4 cm) CSD silencing for four different head models, using HD-EEG simulations. In our simulations, CSD propagation happens in the first 30 minutes and starts on the left hemisphere. Based on the results in Fig. 20, our algorithm detects the CSD propagation correctly.

**Fig. 20:**
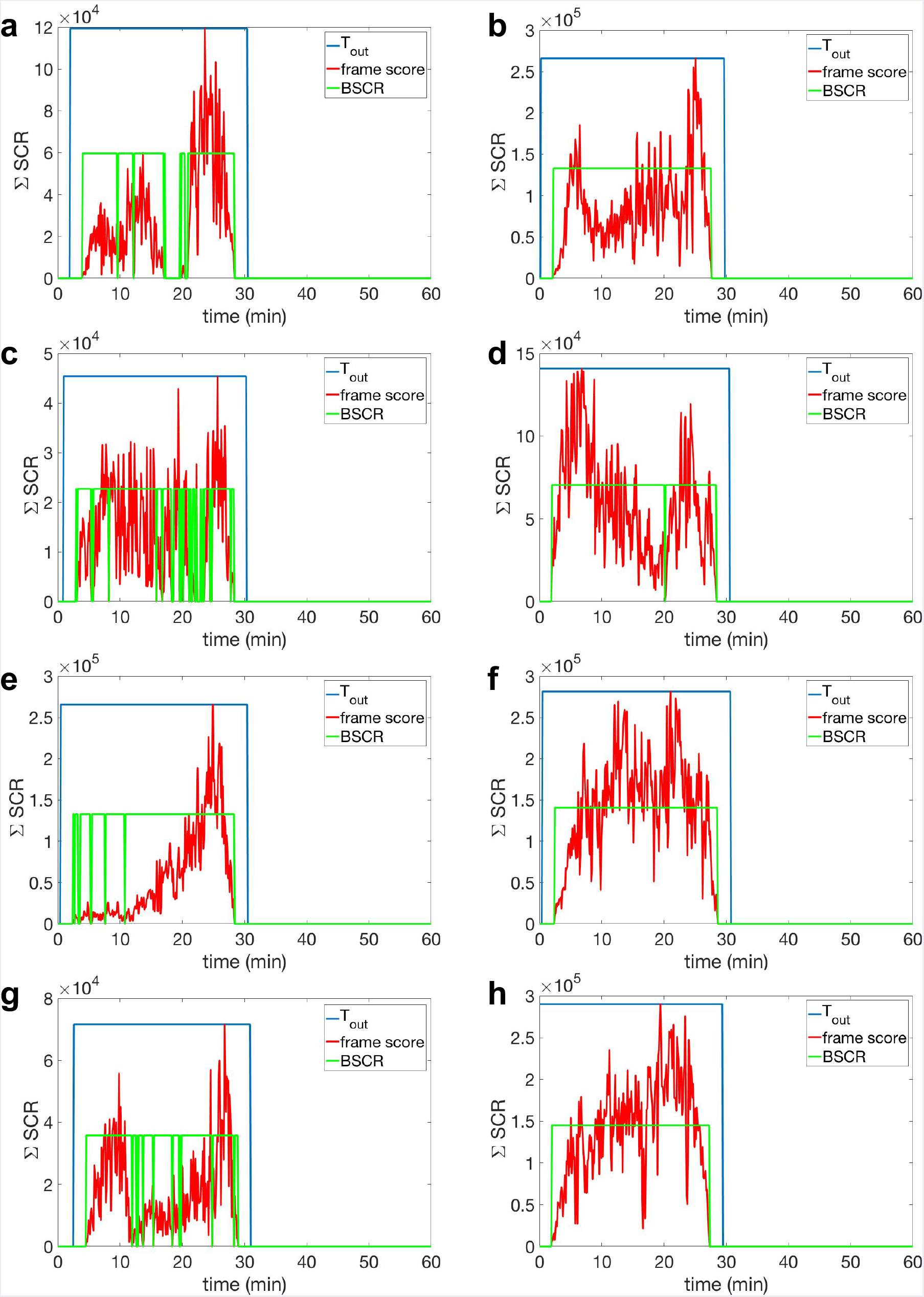
Performance of the proposed algorithm in detection of CSD with different widths of suppression using HD-EEG (*N_ch_* = 340). (a), (c), (e), and (g) are the detection results for narrow suppression width (1 cm wide) and similarly (b), (d), (f), and (h) are for wide suppression width (6.5 cm) for OASIS1, OASIS2, OASIS4, and OASIS5 headmodels respectively. In these simulations CSD propagation happens in the first 30 minutes and the detected time of propagation is given by binary signal, *T_out_*, where non-zero level indicates the presence CSD propagating wave.

Fig. 21a and b, respectively show the detection results for two complex patterns of CSD waves, namely, propagation with semi-planar wavefronts, and on single gyrus. Based on these results, our detection algorithm successfully detects the entire 30 minutes propagation of semi-planar wavefronts of a CSD wave with 2.5 cm width of suppression. However, for a narrow CSD wavefront (5mm wide) propagating on a single gyrus, our detection algorithm misses the first few minutes of propagation due to the small total area of suppression which is below the chosen threshold for the area of bounding boxes (see section III-B).

**Fig. 21:**
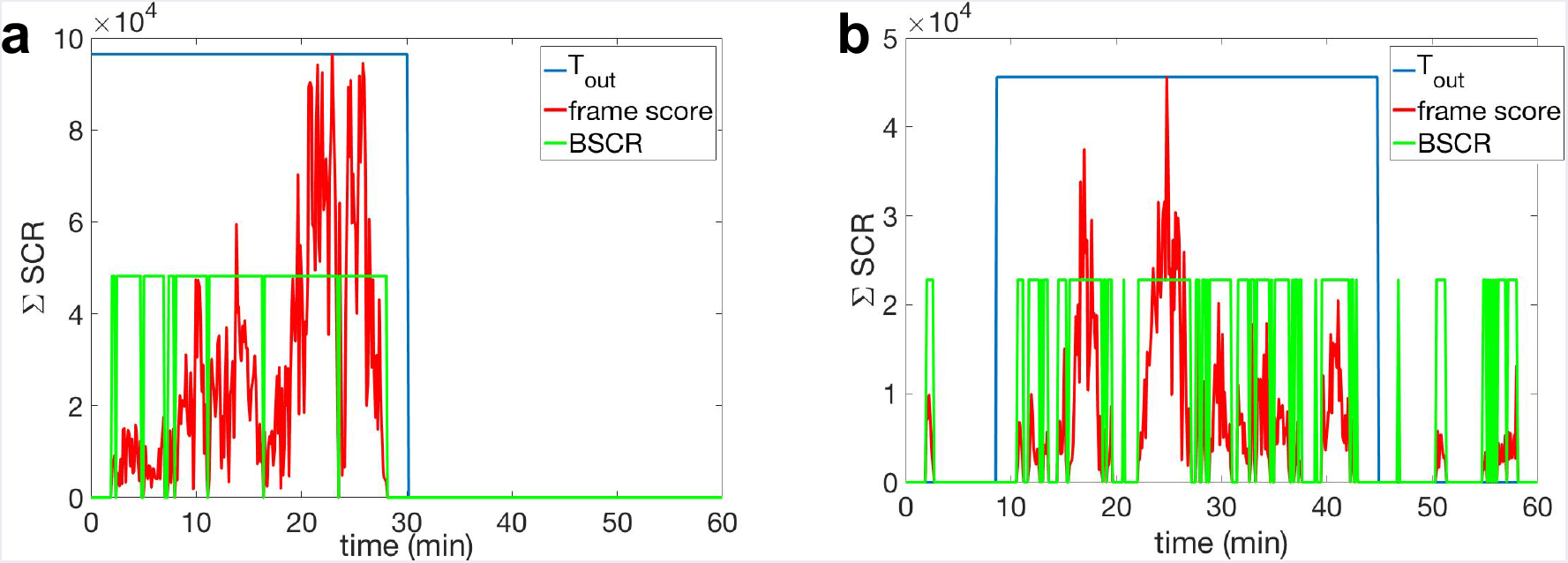
Performance of the proposed algorithm in the detection of complex patterns of CSD propagation using HD-EEG (*N_ch_* = 340) for OASIS2 head model. a) The detection result of CSD propagation with multiple semi-planar wavefronts with 2.5 cm width of suppression (see Fig. 4a). b) The detection result of a narrow CSD wave propagating on a single gyrus, with 5 mm width of suppression (see Fig. 4b). The CSD spread happens in the first 30 minutes for (a) and 45 minutes in (b). The estimated time of propagation is given by the binary signal *T_out_*, whose non-zero level indicates the presence of CSD propagating wave. Note that in (b), our detection algorithm misses the first few minutes of propagation due to the small area of single CSD wavefront.

For a fair comparison, we simulate an hour of EEG signals where there is no propagating electrical suppression, which is correctly rejected by our algorithm (see Fig. 22).

**Fig. 22:**
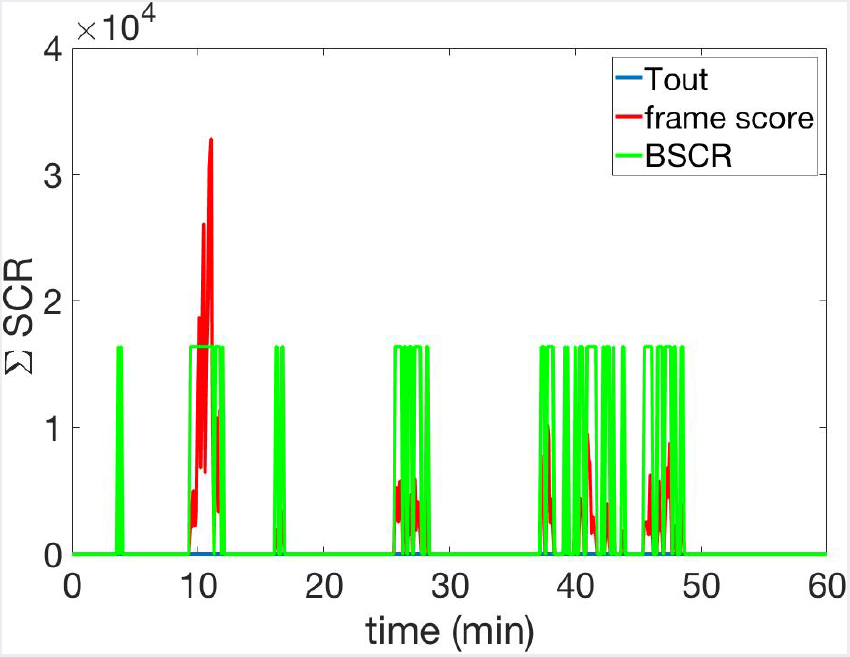
Performance of the proposed detection algorithm in rejecting non-CSD evanescent waves. In a simulation with no CSD propagation, the algorithm correctly outputs *T_out_* as zero during the whole 1-hour simulation.

## IV. DISCUSSIONS AND CONCLUSIONS

Our detection algorithm can detect different types of CSD waves, including narrow and complex patterns of CSD, using HD-EEG as long as the following conditions hold: (i) the CSD wave must have a width of suppression of at least 5 mm; (ii) the wavefronts overall must have a total area of suppression of at least 20 mm^2^. To the best of our knowledge, this threshold is less than the minimum reported area of suppression in the literature which is 80 mm^2^ for a single CSD wave in stroke patients [28]; and (iii) the wave must propagate for at least 5 minutes. Note that some short and temporally isolated CSDs [28] may not be detected using this algorithm.

This detection algorithm is not real-time. The speed of diagnosis and treatment in brain injuries is an important factor and can affect patients’ quality of life after the injury. Future work will extend this algorithm to enable real-time detection.

EEG grids of 340 electrodes might indeed seem unrealistic in today’s clinical settings, especially in situations where EEG electrode placement is limited by skull fractures, wounds, sutures, etc. This poses two engineering challenges. First, on the instrumentation side, when the clinical situation allows, how can one place hundreds of electrodes on the scalp within few minutes? Second, on the signal-processing side, can the algorithms for CSD detection be advanced to detect using limited (nonuniform) EEG arrays, and what would these arrays look like? Even for these lower electrode-count systems, our techniques might still be useful. Specifically, the preprocessing steps of our detection algorithm (see Fig. 15) can enhance the single-channel visual inspection of our simulated EEG signals for CSD detection. Going beyond TBI, migraine CSDs have smaller CSD widths [6], which makes their detection hard. Migraine CSDs have to be detected noninvasively, and thus HD-EEG is likely required.

Anauthor-notes shortcoming of our model is that it does not include low EEG baseline amplitudes, e.g., near lesions and in ischemic tissues. While obtaining realistic models of this aspect difficult, we believe it is important because it will also directly affect our preprocessing step where the cross correlation and thresholding computation might miss some CSDs because the baseline EEG itself is smaller. Simply normalizing each EEG channel (or signals in source space, obtained, e.g., after source localization or Laplacian filtering) can result in blowing up of noise, thus the algorithm will need to be adapted carefully. Owing to these significant complications in this problem, addressing this issue is left as future work. Finally, the algorithm needs to be tested through in-vivo recordings.

## Footnotes

1 The following three sources add up to this number: National Hospital Discharge Survey (NHDS), 2010(https://www.cdc.gov/nchs/nhds/nhds_tables.htm); National Hospital Ambulatory Medical Care Survey (NHAMCS), 2010 (https://www.cdc.gov/nchs/ahcd/web_tables.htm); and, National Vital Statistics System (NVSS), 2010 (https://www.cdc.gov/nchs/nvss/index.htm). All data sources are maintained by the CDC National Center for Health Statistics.

2 National Vital Statistics System (NVSS), 2006–2010. The data source is maintained by the CDC National Center for Health Statistics.

3 http://www.oasis-brains.org

4 For all of the spherical plots in this paper, we use the SSHT package [33] available at: http://www.jasonmcewen.org

5 https://surfer.nmr.mgh.harvard.edu

6 We report results based on suppression widths, so this aspect is not discussed in our results section.

7 This envelope extraction algorithm first detects the tallest peak in the whole signal and ignores the other peaks in the distance of 20 samples (14 seconds) from that. Then it repeats for the remaining peaks and iterates until it considers all of the remaining peaks in the signal. It connects the extracted peaks using spline interpolation to obtain the envelope of the signal.

8 In some relevant works such as [22], these two exchange pump equations are combined.

## ACKNOWLEDGMENTS

We thank the reviewers of this paper whose insightful comments led to significant improvements in the paper. We also thank Shawn Kelly and Marlene Behrmann for helpful discussions. We acknowledge the support of a grant from the Chuck Noll Foundation for Brain Injury Research, and CMU CIT Innovation seed funding.

## APPENDIX

### PDEs in CSD Mesoscale model

This section closely follows Tuckwell’s work in [14] and [22]. Extracellular space (ECS) concentrations of the six components are denoted by 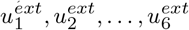 with the order as *K*^+^, *Ca*^++^, *Na*^+^, *Cl*^−^, *T_E_* and *T_I_* respectively, and intracellular space (ICS) concentrations of four components, namely, *K*^+^, *Ca*^++^, *Na*^+^, *Cl*^−^, are denoted by 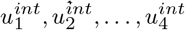, respectively. The dynamics of the ECS concentrations are described by coupled parabolic partial differential equations in (1). ICS concentrations are updated based on the difference between ECS concentrations and their resting equilibrium values, the latter denoted using superscript *R*. For the intracellular concentration of *K*^+^, *Na*^+^, *Cl*^−^ (*i* = 1, 3, 4):

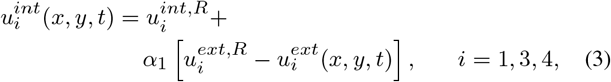

and for updating the *Ca*^++^ concentration (*i* = 2) in ICS:

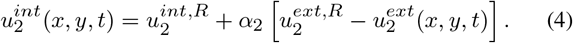

The membrane potential *V_M_* is computed using the Goldman formula [42]:

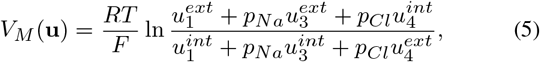

where *p_Na_* and *p_Cl_* are relative permeabilities of sodium and chloride respectively and u is the vector of intra and extracellular concentrations.

The Nernst potential for all of four ions is given by:

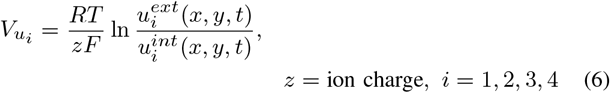

The Nernst equation is based on the energy of chemical reactions, and the ion charge *z* is the number of electrons in such a reaction equation which is related to the amount of transferred charge at the end of the reaction.

In (1), the 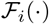 function is the flux term for each component which consists of two main parts, active pump currents and passive fluxes through ion channels. This function makes the PDEs in (1) coupled since it is a function of ECS and ICS concentrations specified by **u**. For *K*^+^ (*i* = 1) this term is calculated as below:

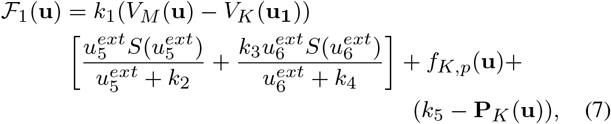

where the *S*(.) function is a step function. In 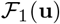 expression, the first term is the passive potassium flux through ligand-gated ion channels which is related to the neurotransmitter concentrations. **P**_*K*_ is called “potassium active pump,” and it determines the strength of the *K*^+^ pump. In addition, *f_K,_p__* is the passive potassium flux through voltage-gated channels defined as below:

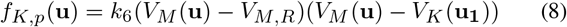

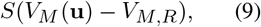

where *V_M,R_* is the resting membrane potential.

We have two separate pump equations for sodium and potassium^8^.

For potassium we have:

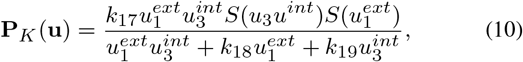

and for sodium:

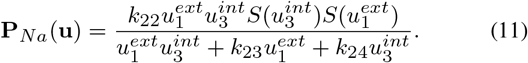

Similarly, for calcium (*i* = 2), the 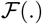 function in (1), is defined as below:

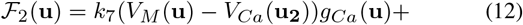

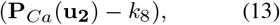

where the new term (as compared to (7)), is the calcium conductance *g_Ca_* (**u**) and is defined as below:

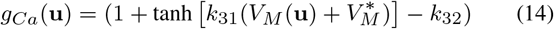

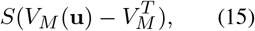

where 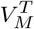 is a cut-off potential and *k*_32_ is defined as below to ensure a smooth rise of *g_Ca_* (**u**) from zero:

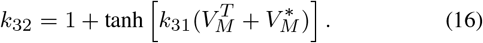

In addition, “calcium active pump” is defined as below:

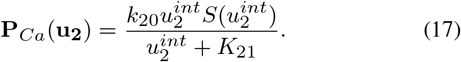

Similar equations exist for sodium and chloride ion fluxes (i=3,4):

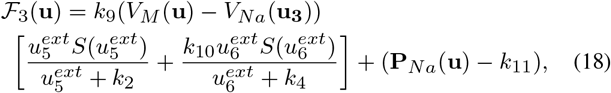

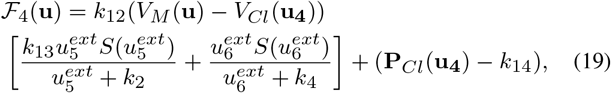

where the “chloride active pump” is defined as below:

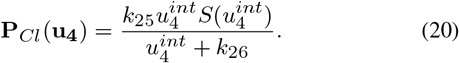

Finally, based on the fact that the rates of transmitter release are proportional to the calcium flux, 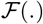 functions of neurotransmitter *T_E_* and *T_I_* are given as:

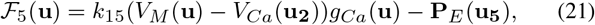

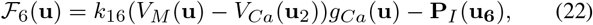

where

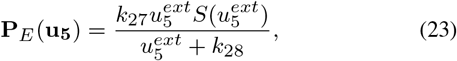

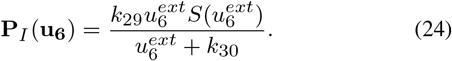

The pumps in (23) and (24) operate to return these neurotransmitters, such as GABA and glutamate, into the glial cells. In this model, the effect of glutamate release from glial cells during CSD propagation is ignored [22].

### KCl stimulation to instigate CSDs

In this model, the CSD propagation is instigated using local elevation of ECS potassium concentration using potassium-chloride (KCl) stimulation with a spatial-exponential profile as below (closely following [22]):

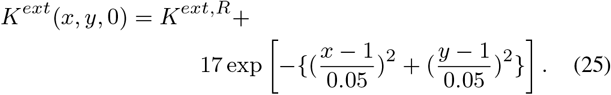

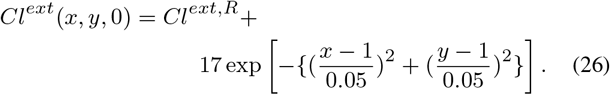

### Using PDEs to simulate CSD waves

To solve the coupled parabolic 2D partial differential equations in (1), they are discretized as below, using Euler’s method (closely following [14]):

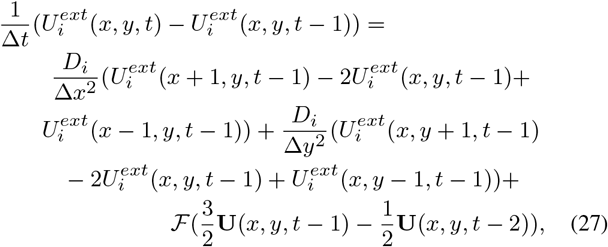

where 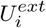 and **U** are the discretized versions of 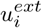 and **u** (ECS concentrations) respectively and *D_i_* is the “scaled” diffusion coefficient of the corresponding component 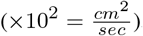. In (27), Tuckwell uses two previous time indices, *t* − 1 and *t* − 2, to make the updates of the parameters smoauthor-notes. The spatial step size of this mesoscale model is 0.78 millimeter (△*x* = △*y*), and the temporal step size (△*t*) is 0.7 second. The dimensions of the 2D plane in this simulation is 15.6 cm×15.6 cm (see Fig. 23a).

**Fig. 23:**
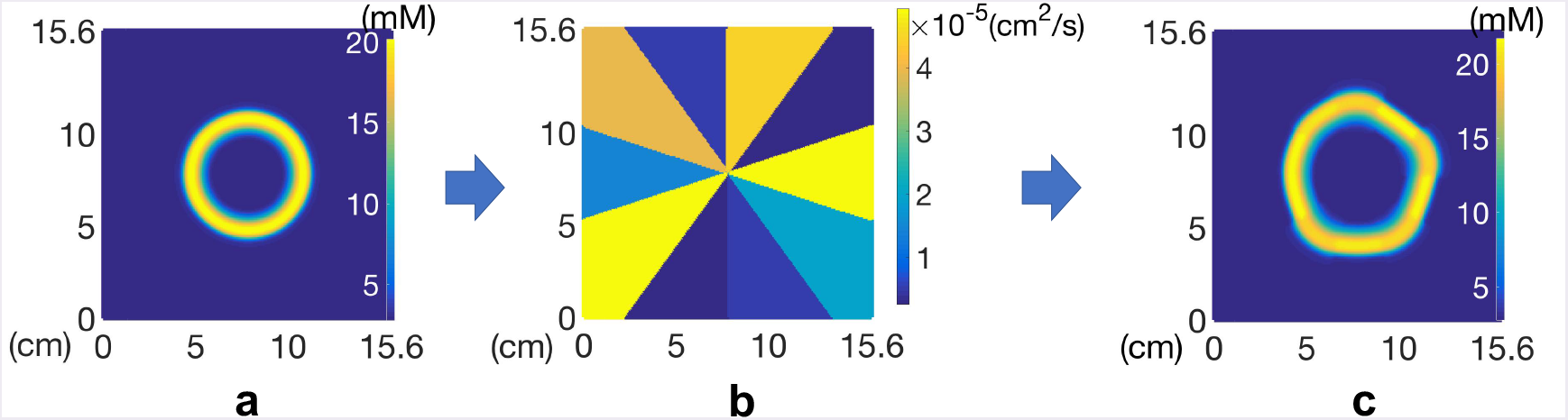
a) Homogeneous model of CSD based on [22], b) spatial profile of potassium diffusion in the range of 0.12 × 10^−5^ to 2.4 × 10^−5^ *cm*^2^ /*s* which is distributed among 10 sectors of the 2D plane, c) the resulting heterogeneous model of CSD propagation after introducing the new spatial distribution of potassium diffusion coefficient (in b) to the Tuckwell’s model.

The initial conditions of equations in (27) are chosen in the way to put the ions’ concentrations at rest (**u**^*R*^) (see [22] for the values):

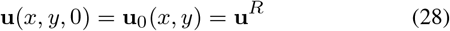

The boundary conditions are fixed at the initial concentrations. Since the propagation starts from the central point of the 2D plane, this assumption is reasonable.

### Heterogeneous model

To take into account the heterogeneity of the ion diffusion coefficients on the CSD waves, as explained in Section II-A, we define 10 sectors (36° each) on this 2D plane and for each sector we arbitrarily assign different diffusion coefficients to potassium (in the range of 5 to 100% of the actual potassium diffusion coefficient 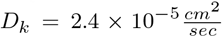 [43]), which is a key component in CSD propagation. The spatial profile of potassium diffusivity is shown in Fig. 23.b. This results in an approximation of heterogeneous ECS for CSD propagation. As it is illustrated in Fig. 23.c, CSD propagates at different speeds, and this causes a deformed ring shape for the CSD wave.

### Upper bound of propagation speed in 2D

As mentioned in Section III, in the spatiotemporal score of OBBoxes, we check the speed of propagation to be less than or equal to 8 mm/min, which is the maximum reported speed for CSD propagation. However, the cylindrical projection (see Section III-A) causes some changes in the propagation speed, i.e., the speed of each wavefront in 2D images (*I_smooth_*) is different from its corresponding speed on the head model. Fig. 24 shows the relation between the wavefront’s speed before and after the cylindrical projection. Without loss of generality, we assume that CSD starts to propagate at the north pole of a head model with radius *r* with an angular speed of *ω* (Fig. 24a). Based on equation 29, the speed of CSD wavefront on the 2D plane, after cylindrical projection, is *rω* sin(*θ*), which is always less than or equal to the maximum reported speed of CSD propagation on head.

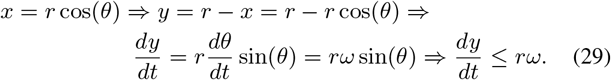

**Fig. 24:**
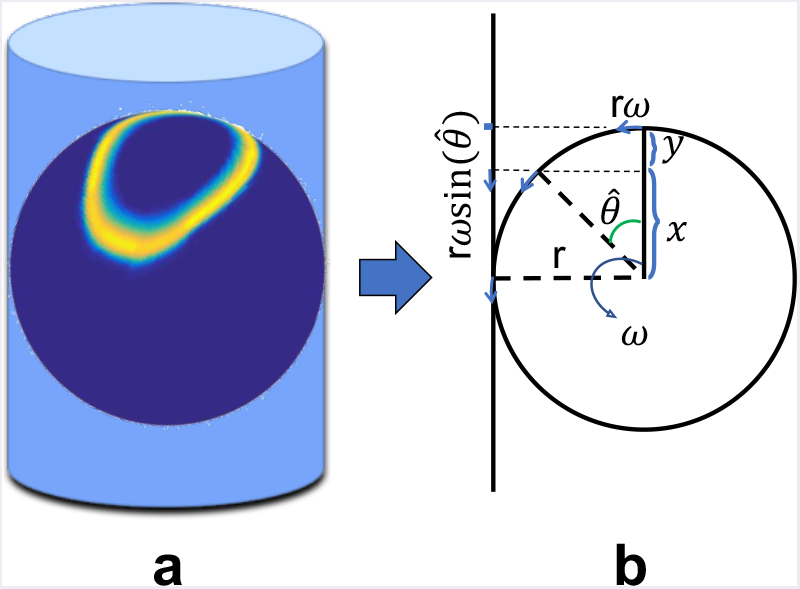
Upper bound limit of propagation speed in 2D: a) In this example, the CSD focal point is at north pole of this sphere, and the cylindrical projection is intentionally oriented in a way to cause maximum possible distortion in the shape of CSD in 2D; b) the mathematical calculation which shows the relation between the angular speed of CSD wavefronts (*ω*) and its speed of propagation on 2D (*rω* sin(*θ*)). Based on this calculation, the speed of CSD on the 2D image (*I_BW_*) is always less than or equal to the maximum reported speed on the head model (8 mm/min).

